# A microbiota-responsive polyfunctional cytotoxic CD4⁺ T-cell state promotes mucosal inflammation in ulcerative colitis

**DOI:** 10.64898/2026.08.10.743953

**Authors:** John P Thomas, Sherine H Kottoor, Jonathan W Lo, Tyler Wooldridge, Hajir Ibraheim, Jonathan Digby-Bell, Nathalie Lambie, Marton Olbei, Balazs Bohar, Charlotte Wong, Eliyas Maroof, Yiwen Cao, Reema Baskar, Matthew Madgwick, Domenico Cozzetto, Hiromi Kudo, Robert Goldin, Nik Matthews, Tamas Korcsmaros, Nick Powell

**Affiliations:** Department of Metabolism, Digestion and Reproduction, Imperial College London, London, United Kingdom; UKRI MRC Laboratory of Medical Sciences, Hammersmith Hospital Campus, London, UK; School of Immunology and Microbial Sciences, King’s College London, London, UK; Centre for Inflammation Biology and Cancer Immunology, King’s College London, UK; NIHR Imperial BRC Genomics Facility, Imperial College London, London, UK; Department of Inflammatory Bowel Disease, St Mark’s National Bowel Hospital, London, UK; Genome Institute of Singapore, A*STAR, Singapore; Gut Microbes and Health Programme, Quadram Institute Bioscience, Norwich Research Park, Norwich, UK; Earlham Institute, Norwich Research Park, Norwich, UK

## Abstract

Ulcerative colitis (UC) is characterised by chronic colonic inflammation with marked heterogeneity in disease severity and therapeutic outcomes. Here, we define a spatially organised, polyfunctional cytotoxic CD4⁺ T-cell state associated with mucosal inflammation and adverse therapeutic outcomes in UC. Integrating *ex vivo* T-cell receptor stimulation with multi-cohort bulk and single-cell transcriptomics and multiparameter flow cytometry, we show that GZMB⁺ CD4⁺ T cells are preferentially enriched in inflamed UC mucosa, but not peripheral blood, and co-express cytotoxic molecules, Th1- and Th17-associated cytokines and chemokines, and immunoregulatory receptors. Single-cell analyses implicate inflammatory cytokine and antigen-presentation signals in the acquisition or maintenance of this state. High-resolution spatial profiling localised this programme predominantly to Th17 cells, which were preferentially enriched within multicellular inflammatory and tertiary lymphoid structure-associated niches. Across independent patient cohorts, a transcriptional signature derived from this state increased with endoscopic disease severity and was associated with reduced response to anti-TNF and anti-IL-12/23p40 therapies. Adoptive transfer of *Gzma/Gzmb*-deficient rather than wild-type CD4⁺ T cells into *Rag2*-deficient recipient mice markedly attenuated experimental colitis and abrogated the polyfunctional cytokine phenotype, demonstrating that granzyme-dependent effector activity is a key mechanism driving CD4^+^ T-cell-mediated intestinal inflammation. Finally, human host-microbiome analysis linked this programme to intestinal dysbiosis, while transfer of dysbiotic microbiota promoted the emergence of a corresponding state *in vivo*. Collectively, these findings define a microbiota-responsive, spatially organised polyfunctional cytotoxic CD4⁺ T-cell programme that contributes to intestinal inflammation and is associated with disease severity and treatment resistance in UC.

## INTRODUCTION

Ulcerative colitis (UC) is the most common form of inflammatory bowel disease (IBD), characterised by relapsing and remitting chronic inflammation of the colonic mucosa^1,2^. It is marked by substantial inter-individual heterogeneity in terms of disease severity, treatment response, and long-term clinical outcomes. Despite major advances in the therapeutic armamentarium in UC, including advanced therapies targeting cytokine signalling (e.g. anti-TNF, anti-IL12/23p40) or lymphocyte trafficking pathways (e.g. anti-α4β7 integrin), most patients (60-70%) fail to achieve sustained clinical and endoscopic remission^2^. These clinical challenges highlight our incomplete understanding of the immune mechanisms that sustain chronic intestinal inflammation and underpin therapeutic resistance in UC.

Historically, UC had been regarded as a CD4⁺ T-helper 2 (Th2) driven disease^3^. However, accumulating evidence indicates that intestinal T-cell immune responses in UC extend beyond Th2 T-cell programmes, encompassing dysregulated Th1 and Th17 cells, as well as unconventional T-cell populations such as gamma delta (γδ) T cells, invariant natural killer T (iNKT) cells, and mucosal-associated invariant T (MAIT) cells^4,5^. Despite these advances, defining pathogenic T-cell states relevant to human disease remains challenging. Most studies rely on static immune profiling approaches that provide limited insight into the functional immune programmes that emerge within chronically inflamed tissues. Moreover, the spatial organisation of mucosal immune responses and its contribution to disease progression and therapeutic outcomes in UC remains incompletely understood despite emerging insights^6,7^. Whether discrete functional T-cell states underlie clinical heterogeneity and treatment resistance also remains unclear. Addressing these questions requires integrative approaches that combine functional immune interrogation with high-resolution molecular and spatial profiling directly in patient tissues.

To address these gaps, here we combined *ex vivo* T-cell receptor (TCR) stimulation of patient-derived colonic lamina propria mononuclear cells (LPMCs) with multimodal analyses spanning multi-cohort bulk and single-cell tissue transcriptomic datasets, multiparameter flow cytometry, high-resolution spatial transcriptomics, and data from independent clinical trial cohorts. This integrative framework revealed a previously unrecognised spatially organised, polyfunctional cytotoxic CD4⁺ T-cell state in UC. We demonstrate that this programme associates with disease severity, treatment resistance to multiple biologics, and gut microbial dysbiosis. Importantly, genetic ablation of cytotoxic effector function in CD4^+^ T-cells attenuates experimental colitis, establishing this programme as functionally required for colitis development. Collectively, these findings unveil a distinct polyfunctional cytotoxic CD4⁺ T-cell state as a key feature of UC pathogenesis, providing novel insights into the mechanisms underpinning disease severity and therapeutic resistance.

## RESULTS

### 1. *Ex vivo* TCR activation of UC patient-derived colonic LPMCs reveals induction of cytotoxic effector programmes

To define inducible mucosal T-cell responses in UC, colonic LPMCs isolated from patients with moderate-to-severe active UC were stimulated *ex vivo* using anti-CD3 agonistic antibodies to activate TCR signalling (Figure 1A). Transcriptomic analysis of stimulated versus unstimulated LPMCs identified 693 differentially expressed genes (DEGs) (absolute log2FC >1, adjusted p value < 0.05) (Figure 1B). These included marked upregulation of genes encoding cytokines (e.g. *IL22, IFNG, IL17A, TNF, IL2*), cytokine receptors (e.g*. IL23R, IL12RB2, IL21R*), chemokines (e.g. *CXCL10, CCL8, CXCL9, CXCL11*), immune regulatory molecules (e.g. *TNFRSF9, CTLA4, PDCD1, LAG3*), and cytotoxic effector molecules (*CRTAM, FASLG, GZMB, PRF1, GZMH,* and *NKG7*) (Figure 1C).

**Figure 1:**
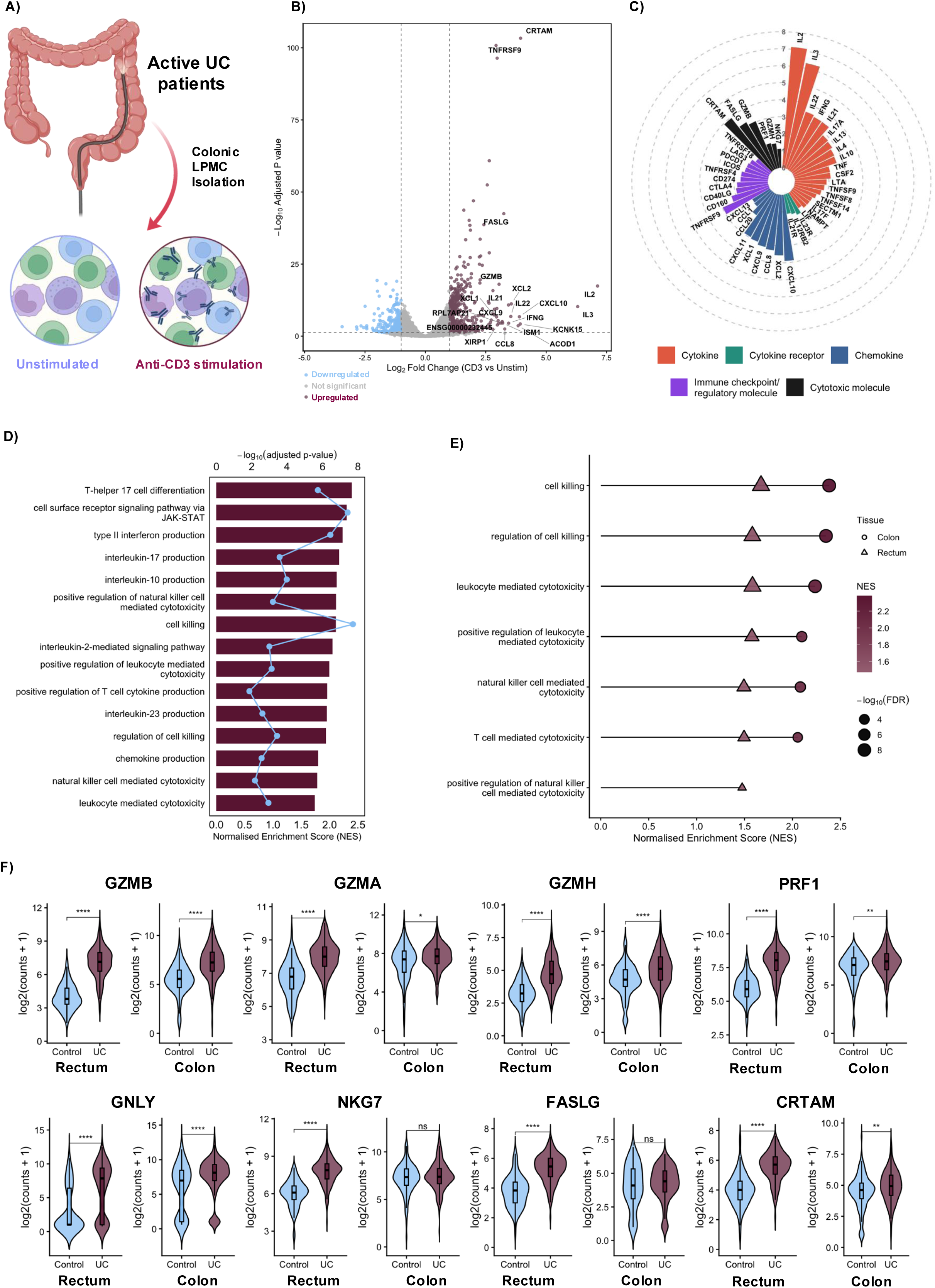
T-cell receptor activation induces cytotoxic effector programmes in UC colonic mucosa. **A,** Schematic of the experimental design. Colonic lamina propria mononuclear cells (LPMCs) isolated from four patients with active UC were divided into paired unstimulated and anti-CD3-stimulated conditions before transcriptomic profiling. **B,** Volcano plot showing differential gene expression in anti-CD3 stimulated LMPMs vs unstimulated LPMCs. Dashed lines indicate the thresholds used to define differentially expressed genes (absolute log₂ fold change >1 and Benjamini-Hochberg-adjusted *P* < 0.05). Selected inflammatory and cytotoxic genes are labelled **C,** Circular bar plot of selected genes upregulated following anti-CD3 stimulation, grouped by functional category. Bar length represents log₂ fold change relative to unstimulated LPMCs. **D,** Gene set enrichment analysis of upregulated Gen Ontology (GO) Biological Process terms following anti-CD3 stimulation. Bars show normalised enrichment scores (NES), while points show −log₁₀(adjusted *P* value). **E,** Enrichment of cytotoxicity-associated GO pathways in UC compared to non-IBD colonic and rectal tissues within the IBD TaMMA resource. Symbol shape: tissue, colour: NES, symbol size: −log₁₀(adusted *P* value). **F,** Expression of selected cytotoxic effector genes in rectal and colonic tissues from patients with UC and non-IBD controls within the TaMMA resource. Violin plots show sample-level log₂(counts + 1) distributions. Groups were compared separately within each tissue using two-sided Mann–Whitney *U* tests. ns, not significant; \**P* < 0.05; \*\**P* < 0.01; \*\*\**P* < 0.001; \*\*\*\**P* < 0.0001. FDR, false-discovery rate; LPMC, lamina propria mononuclear cell; TaMMA, IBD Transcriptome and Metatranscriptome Meta-Analysis; UC, ulcerative colitis.

Gene set enrichment analysis (GSEA) identified multiple upregulated pathways in stimulated colonic LPMCs. Cytotoxic effector pathways, including cell killing and leukocyte-mediated cytotoxicity, dominated the top 15 upregulated pathways, alongside inflammatory cytokine signalling and T-cell activation programmes such as Th17 differentiation, JAK-STAT signalling, and production of IFN-γ, IL-17 and IL-10 (Figure 1D, Supplementary Figure 1).

Given the emergence of cytotoxic pathways following TCR activation in patient-derived colonic LPMCs, we next sought to evaluate whether these findings were reproducible across independent UC patient cohorts. To do this, we utilised the IBD Transcriptome and Metatranscriptome Meta-Analysis (TaMMA) resource^8^, comprising bulk transcriptomics datasets of IBD patient-derived and control tissues from 26 independent studies. Cytotoxic pathways including leucocyte-mediated cytotoxicity, T-cell-mediated cytotoxicity, and cell killing, were significantly enriched in both rectal and colonic tissue samples from UC patients compared with non-IBD controls (Figure 1E). At gene level, multiple cytotoxic effector genes, including *GZMB*, *GZMA*, *GZMH*, *PRF1*, *GNLY*, *NKG7*, *FASLG*, and *CRTAM*, were significantly increased (Figure 1F).

Collectively, these findings identify inducible cytotoxic gene programmes as a conserved feature of activated UC mucosal immune cell responses that are enriched across multiple independent patient cohorts.

### 2. CD4⁺ T cells are a major source of cytotoxic effector activity and display a polyfunctional inflammatory phenotype in the UC colonic mucosa

To identify the cellular sources of cytotoxic effector programmes in UC, we performed multiparameter flow cytometry on colonic LPMCs from patients with endoscopically active disease (Ulcerative Colitis Endoscopic Index of Severity (UCEIS) score ≥ 2, N = 32) and non-IBD controls (N = 17) (Supplementary Figure 2A) profiling expression of the cytotoxic effector molecule GZMB, alongside key inflammatory cytokines (IFN-γ, TNF-α, IL-22 and IL-17A), across conventional CD4^+^ and CD8^+^ T cells and unconventional T-cell subsets (γ/δ T cells, iNKT cells, and MAIT cells (Figure 2A, Supplementary Table 1).

**Figure 2:**
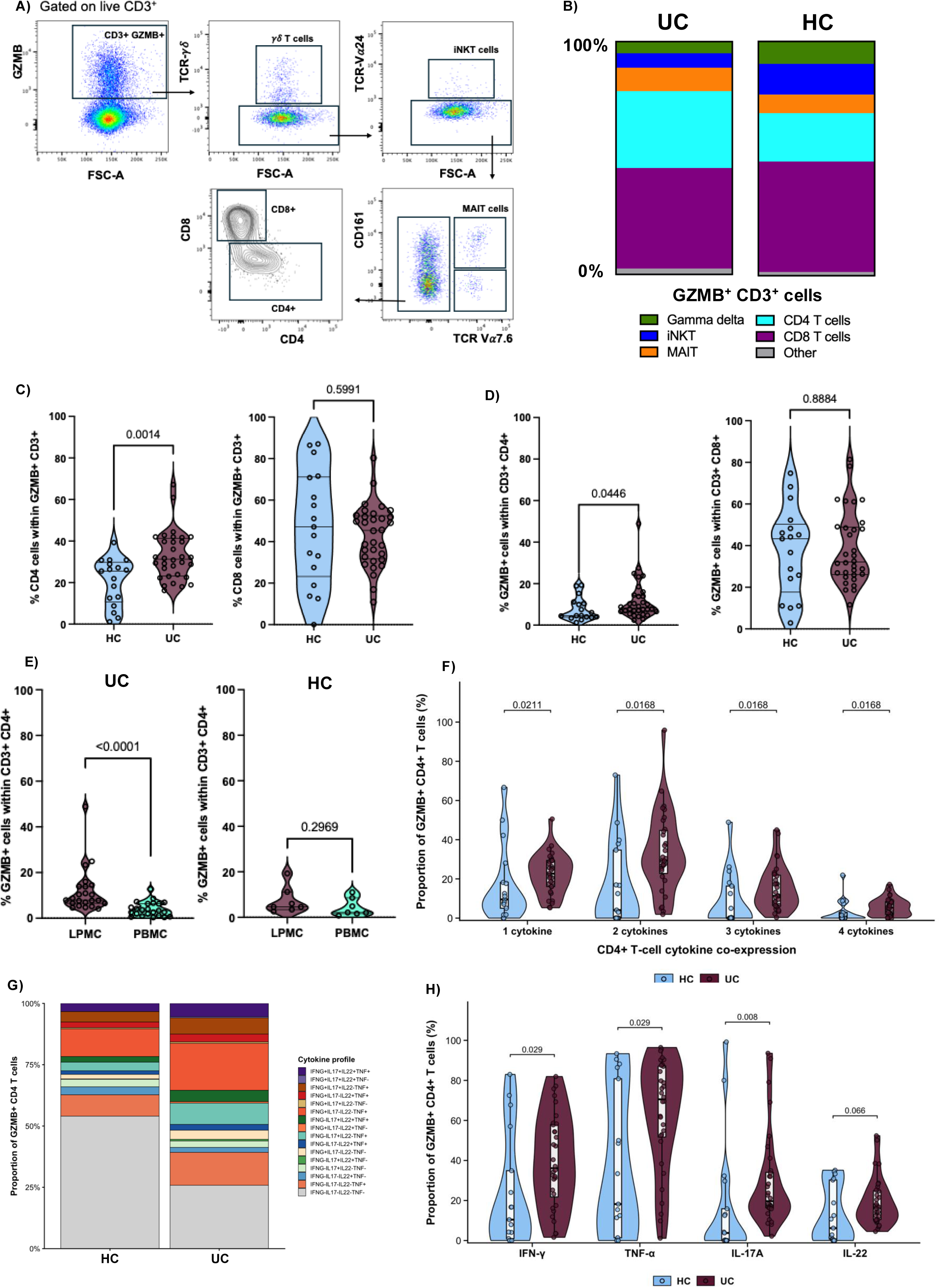
GZMB⁺ CD4⁺ T cells are preferentially enriched and exhibit a polyfunctional cytokine profile in UC colonic mucosa. **A,** Representative flow cytometry gating strategy used to identify GZMB-expressing T-cell populations among live CD3⁺ lymphocytes. **B,** Mean relative composition of GZMB⁺CD3⁺ LPMCs in UC patients and non-IBD controls. **C,** Proportions of GZMB⁺CD3⁺ T cells belonging to CD4⁺ or CD8⁺ lineages in UC (n = 32) and non-IBD controls (n = 17). **D,** Proportions of GZMB⁺ cells within the CD3⁺CD4⁺ and CD3⁺CD8⁺ T-cell populations. **E,** Proportions of GZMB⁺ cells within paired mucosal CD3⁺CD4⁺ LPMCs and circulating CD3⁺CD4⁺ PBMCs from patients with UC (n = 24) and non-IBD controls (n = 7). **F,** Proportions of mucosal GZMB⁺ CD4⁺ T cells expressing one, two, three or all four measured cytokines: IFN-γ, TNF-α, IL-17A and IL-22. **G,** Mean relative composition of the 16 mutually exclusive cytokine expression profiles within mucosal GZMB⁺ CD4⁺ T cells from UC and non-IBD tissues. **H,** Proportions of mucosal GZMB⁺ CD4⁺ T cells expressing each cytokine irrespective of co-expression with the other cytokines. In **C**, **D**, **F** and **H**, each point represents one participant; violin plots show the distribution and embedded box plots denote the median and interquartile range. Independent groups were compared using two-sided Mann-Whitney U tests. Paired LPMC-PBMC measurements in **E** were compared using two-sided Wilcoxon signed-rank tests. In **F** and **H**, *P* values were adjusted using the Benjamini-Hochberg procedure. HC, non-IBD control; LPMC, lamina propria mononuclear cell; MAIT, mucosal-associated invariant T cell; PBMC, peripheral blood mononuclear cell; UC, ulcerative colitis.

Conventional T-cells accounted for the majority of GZMB⁺ T-lymphocytes across all samples (Figure 2B). CD8⁺ T cells represented the largest component of the GZMB⁺CD3⁺ compartment in UC and non-IBD controls (median 45.4% [IQR 31.6–52.4%] versus 47.1% [IQR 23.2–71.2%], respectively; p = 0.60, Mann-Whitney U test) (Figure 2C). In contrast, CD4⁺ T cells constituted a greater proportion of GZMB⁺CD3⁺ cells in UC compared to controls (31.3%, IQR 23.2-41.2% vs. 25.6%, IQR 10.6-29.9%; p = 0.0014), thereby establishing CD4⁺ T cells as a major disease-enriched source of mucosal GZMB.

A complementary analysis examined the frequency of GZMB expression within CD4^+^ and CD8^+^ T cells (Figure 2D, Supplementary Figure 2B). The proportion of GZMB⁺ cells within the CD4⁺ T-cell population was significantly higher in UC than in non-IBD control (8.3%, IQR 6.5-14.3% vs. 4.6%, IQR 3.8-10.8%; p = 0.045), whereas the corresponding proportion within CD8⁺ T cells did not differ between groups (32.1%, IQR 26.0-49.0% vs. 43.0%, IQR 17.6-50.3%; p = 0.89). These findings further support the UC-specific enrichment of GZMB-expressing CD4⁺ T cells.

To determine whether this GZMB⁺ CD4⁺ enrichment extended systemically, we compared paired mucosal and circulating compartments (UC, N = 24; non-IBD controls, N = 7). CD8⁺ T cells dominated the circulating GZMB⁺CD3⁺ compartment in both groups, (non-IBD controls, 57.8%; UC, 47.8%), with CD4⁺ T cells contributing a smaller fraction (non-IBD controls, 18.6%; UC, 19.8%; Supplementary Figure 2E). Unlike the mucosal compartment, neither CD4⁺ GZMB⁺ frequency nor CD4⁺ contribution to the circulating GZMB⁺CD3⁺ pool differed significantly between groups (p = 0.99 and p = 0.55 respectively; Supplementary Figure 2F). CD8⁺ GZMB⁺ frequency was significantly increased in circulating UC samples (p < 0.0001), although CD8⁺ contribution to the GZMB⁺CD3⁺ pool was unchanged (p = 0.3407) (Supplementary Figure 2G).

Comparison of the mucosal and circulating compartments showed that among UC samples, CD4⁺ T cells constituted a significantly greater proportion of GZMB⁺CD3⁺ cells in LPMCs than in PBMCs (p < 0.001), and the frequency of GZMB⁺ cells within the CD3⁺CD4⁺ population was also higher in LPMCs (p < 0.0001) (Figure 2E). No significant differences were observed in non-IBD controls. Collectively, these findings identify CD4⁺ T cells as a major source of mucosal GZMB in UC and demonstrate preferential enrichment of this population within inflamed colonic tissue rather than a generalised expansion in the peripheral circulation.

Notably, mucosal GZMB⁺ CD4⁺ T cells in UC demonstrated a polyfunctional cytokine profile relative to non-IBD controls. UC samples contained significantly higher proportions of GZMB⁺ CD4⁺ T cells expressing one cytokine (UC: median 22.7%, IQR 16.1-29.4%; controls: median 9.5%, IQR 5.0-18.2%; adjusted p = 0.0211), two cytokines (UC: median 29.3%, IQR 22.8-44.7%; controls: median 4.1%, IQR 0–34.7%; adjusted p = 0.0168), three cytokines (UC: median 11.3%, IQR 6.4-21.9%; controls: median 0%, IQR 0-16.3%; adjusted p = 0.0168) and four cytokines (UC: median 5.3%, IQR 1.9-8.4%; controls: median 0%, IQR 0-2.9%; adjusted p = 0.0168; two-sided Mann-Whitney U tests with Benjamini-Hochberg correction) (Figure 2F).

Analysis of cytokine co-expression patterns provided further evidence of inflammatory cytokine heterogeneity among GZMB⁺ CD4⁺ T cells in the UC colonic mucosa (Figure 2G and Supplementary Figure 2C). Individual cytokine combinations were compared between UC and non-IBD controls across the 16 mutually exclusive cytokine-expression patterns (Supplementary Figure 2D). Cytokine-negative cells were significantly lower in UC compared to controls (25.9% vs 54.0% respectively, adjusted p = 0.0411). Conversely, several cytokine-positive populations were more highly represented in UC, including IFN-γ⁺ TNF-α⁺ double-positive cells (19.2% vs 11.3% respectively, adjusted p = 0.017) and IL-17A⁺TNF-α⁺ double-positive cells (8.7% vs 3.5% respectively, adjusted p = 0.008). All four triple-cytokine-positive phenotypes were also significantly enriched in UC, including IFN-γ⁺IL-17A⁺TNF-α⁺ cells (6.6% versus 4.3%; adjusted p = 0.0109), IL-17A⁺IL-22⁺TNF-α⁺ cells (4.6% versus 2.3%; adjusted p = 0.0109), IFN-γ⁺IL-22⁺TNF-α⁺ cells (3.2% versus 2.3%; adjusted p = 0.0187) and IFN-γ⁺IL-17A⁺IL-22⁺ cells (0.6% versus 0.5%; adjusted p = 0.0124). Cells expressing all four cytokines were increased in UC (5.4% versus 3.3%; adjusted p = 0.0185). Overall, 14 of the 15 cytokine-positive combinations were significantly enriched in UC.

To determine the contribution of each cytokine to this broader polyfunctional phenotype, cytokine-positive populations were then aggregated irrespective of co-expression with the other cytokines (Figure 2H). GZMB⁺ CD4⁺ T cells expressing IFN-γ, TNF-α and IL-17A were significantly more highly represented in UC than in non-IBD controls (adjusted p = 0.029, adjusted p = 0.029 and adjusted p = 0.008, respectively), whereas the increase in IL-22 expression in UC did not reach the threshold of statistical significance (adjusted p = 0.066; two-sided Mann-Whitney U tests with Benjamini-Hochberg correction). These results suggest that cytotoxic CD4⁺ T cells in UC exhibit increased Th1- and Th17-associated cytokine expression consistent with a polyfunctional effector phenotype.

Taken together, these findings reveal a GZMB⁺ CD4⁺ T-cell state preferentially enriched in inflamed UC colonic mucosa characterised by combined cytotoxic and multi-cytokine effector functions.

### 3. A transcriptionally distinct polyfunctional cytotoxic CD4⁺ T-cell state defines inflamed UC colonic tissue

To determine whether GZMB^+^ CD4⁺ T cells represent a distinct transcriptional state we interrogated single-cell RNA-sequencing data from inflamed UC colonic tissues within the single-cell IBD (scIBD) resource^9^ (Supplementary Figures 3A and 3B). Compared with GZMB^−^ CD4⁺ T cells, GZMB⁺ CD4⁺ T cells displayed coordinated upregulation (log_2_FC > 1, adjusted p <0.05) of cytotoxic effector genes (*GZMA, GNLY, NKG7, PRF1, FASLG*), inflammatory cytokines and chemokines (*IFNG, IL26, CCL4, CCL5, CCR5, CCL20, CXCR6*), and immune regulatory markers associated with activation and exhaustion (*LAG3, HAVCR2, TNFRSF18*) (Figure 3A). The 20 most significantly upregulated genes were used to define a cytotoxic CD4⁺ T-cell gene signature for downstream analyses. GSEA confirmed this transcriptional programme at pathway level. GZMB⁺ CD4⁺ T cells were upregulated for cytotoxicity-associated pathways (cell killing, leukocyte-mediated cytotoxicity, pyroptosis) alongside Th1/IFNg responses, IL-17 production, cytokine and NF-κB signalling, and lymphocyte/macrophage chemotaxis (Figure 3B), mirroring the IFN-γ and IL-17A co-expression identified by flow cytometry.

**Figure 3:**
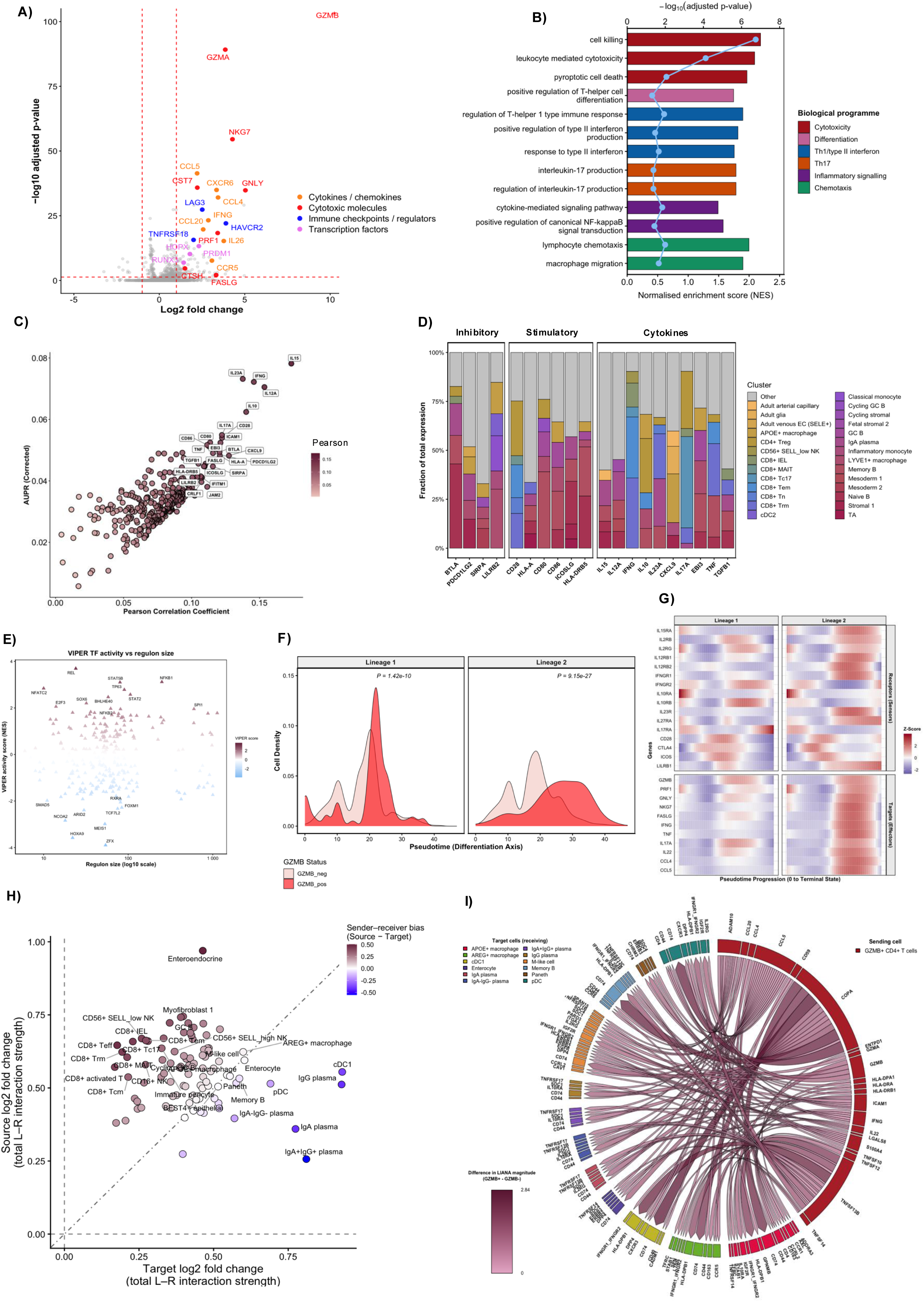
GZMB⁺ CD4⁺ T cells exhibit a transcriptionally distinct, polyfunctional cytotoxic effector state in inflamed UC mucosa. **A,** Differential gene-expression analysis comparing GZMB⁺ and GZMB⁻ CD4⁺ T cells from inflamed UC colonic tissues in the scIBD resource. Dashed lines indicate thresholds of absolute log₂ fold change >1 and Benjamini-Hochberg-adjusted *P* < 0.05. Selected genes are labelled and coloured by functional category. **B,** Gene set enrichment analysis of selected GO Biological Process terms associated with the GZMB⁺ CD4⁺ T-cell state. Bars show normalised enrichment scores (NES), points show −log₁₀(adjusted *P* value), and bar colours denote broader biological programmes. **C,** NicheNet prioritisation of candidate upstream ligands associated with the GZMB⁺ CD4⁺ T-cell transcriptional programme. Ligands are positioned according to their Pearson correlation coefficient and corrected area under the precision-recall curve (AUPR); the top 25 highly ranked ligands are labelled. **D,** Distribution of expression of prioritised inhibitory, stimulatory and cytokine ligands across candidate sender-cell populations in inflamed UC tissue from scIBD. Each bar shows the fraction of total ligand expression attributable to the indicated cellular populations. **E,** VIPER-inferred transcription factor activity in GZMB⁺ relative to GZMB⁻ CD4⁺ T cells. Colour denotes the direction and magnitude of inferred activity. **F,** Density distributions of GZMB⁺ and GZMB⁻ CD4⁺ T cells along two Slingshot-inferred differentiation trajectories originating from CD4⁺ memory T cells. *P* values were calculated using one-sided Wilcoxon rank-sum tests evaluating whether GZMB⁺ cells occupied later pseudotime positions. **G,** Pseudotime-resolved expression dynamics of genes associated with cytokine responsiveness, immune regulation, cytotoxic and inflammatory effector function along the two inferred lineages. Expression was modelled using LOESS smoothing across 50 equally spaced pseudotime positions and standardised to gene-level z-scores; pseudotime progresses from left to right towards the inferred terminal state. **H,** Differential aggregate ligand-receptor communication involving GZMB⁺ relative to GZMB⁻ CD4⁺ T cells, inferred using LIANA. Each point represents a partner population. The y-axis shows the relative fold change in outgoing communication from GZMB⁺CD4⁺ T-cells versus GZMB⁻ CD4⁺ T cells, while the x-axis shows the relative fold change in incoming communication towards GZMB⁺CD4⁺ T-cells versus GZMB⁻ CD4⁺ T cells. Point colour denotes the bias towards outgoing or incoming communication. **I,** Chord diagram showing the top preferential outgoing ligand-receptor interactions from GZMB⁺ CD4⁺ T cells towards lymphoid, myeloid, dendritic and epithelial target populations. Ribbons connect ligands expressed by GZMB⁺ CD4⁺ T cells with their predicted receptors in target populations, with ribbon shading representing the increase in LIANA interaction magnitude relative to GZMB⁻ CD4⁺ T cells. LIANA interactions with CellPhoneDB-derived *P* < 0.05 were retained. AUPR, area under the precision-recall curve; GC, germinal centre; GSEA, gene set enrichment analysis; NES, normalised enrichment score; TF, transcription factor; UC, ulcerative colitis.

To explore upstream signals associated with the acquisition or maintenance of this state, we next applied NicheNet^10^ analysis to this dataset which highlighted cytokines including, IL-15, IL-23A, IFN-γ, EBI3, IL-12A, IL-17A, TNF and IL-10 as the most strongly predicted upstream ligands (Figure 3C). Additional prioritised interactions involved co-stimulatory and co-inhibitory immune-regulatory molecules, including CD28, CD80, CD86, HLA-DRB5, ICOSLG and BTLA, consistent with antigen presentation and chronic immune stimulation. The predicted ligand-target regulatory matrix demonstrated that these prioritised ligands were capable of explaining multiple components of the GZMB⁺ CD4⁺ T-cell transcriptional programme, including genes associated with cytokine responsiveness, immune regulation and cytotoxic or inflammatory effector function (Supplementary Figure 3C). Mapping the expression of these ligands across candidate sender populations in inflamed relative to reference tissue demonstrated that the predicted signals arose from multiple lymphoid, myeloid and epithelial compartments rather than a single cellular source (Supplementary Figure 3D). The major cellular sources of these ligands included memory B cells (*IL27, TNF, IL23A, CD80, CD86, HLA-DRB5*), plasma cells (*IL23A, IL12A, IL15*), germinal cell B cells (*ICOSLG, CD86, EBI3*), naïve B cells (*HLA-DRB5, ICOSLG, BTLA, IL15*), regulatory T cells (*IL10, IL17A, CD28*), CD8^+^ tissue resident and effector memory cells (*IFNG, CD28*), and APOE^+^ macrophages (*CXCL9, CD86*).

Upstream gene regulatory network analysis^11^ further revealed NF-κB components (REL, NF-κB1, and NF-κB2), STAT5B, STAT2, and BHLHE40 as potential transcriptional drivers of this cytotoxic CD4+ T-cell programme, accompanied by reduced activity of transcription factors including HOXA9, MEIS1, ZFX, and ARID2. To delineate the developmental trajectory underlying acquisition of this cytotoxic CD4⁺ T-cell state, we performed pseudotime analysis using the largest single-cell UC dataset^12^ within scIBD. Pseudotime analysis indicated that GZMB⁺ CD4⁺ T cells may emerge from CD4⁺ memory T cells along two differentiation trajectories (Supplementary Figure 3E). These cells occupied a more advanced pseudotime position relative to GZMB⁻ CD4⁺ T cells (Figure 3F), consistent with acquisition of a late effector phenotype. This was supported by increased expression of transcriptional regulators involved in effector T-cell differentiation including *PRDM1*, *RUNX3*, and *HOPX* in the earlier differential expression analysis (Figure 3A). Pseudotime-resolved expression dynamics further demonstrated the coordinated acquisition of immunoregulatory, cytokine-responsive and polyfunctional effector programmes, most prominently along the second differentiation trajectory (Figure 3G). Progression towards the terminal state was accompanied by increased expression of receptors associated with IL-15/IL-2, IL-12, IFN-γ, IL-23 and IL-27 signalling, alongside induction of cytotoxic effector genes, inflammatory cytokines and chemokines. Collectively, these findings support the progressive acquisition of a cytokine-responsive, polyfunctional cytotoxic effector state during CD4⁺ T-cell differentiation.

We next employed ligand-receptor analysis using LIANA^13^ to understand differences in intercellular communication between GZMB⁺ versus GZMB^−^ CD4⁺ T cells. Relative to GZMB⁻ CD4⁺ T cells, GZMB⁺ CD4⁺ T cells exhibited preferentially stronger outgoing communication with B-cell and dendritic-cell populations, while receiving preferentially stronger incoming signals from CD8^+^ T-cells, NK cells and germinal centre (GC) B cells (Figure 3H). Comparably increased bidirectional communication was observed with macrophage populations and epithelial cell types, including enterocytes, Paneth cells and M-like cells.

To resolve the ligand-receptor interactions underlying these cellular communication patterns, we visualised interactions restricted to populations exhibiting preferential incoming, outgoing or bidirectional communication (Figure 3I). Relative to GZMB⁻ CD4⁺ T cells, GZMB⁺ CD4⁺ T cells exhibited higher inferred outgoing ligand-receptor interaction magnitudes across a diverse range of effector pathways, including inflammatory cytokines such as IFN-γ and IL-22, chemokines including CCL20, CCL4 and CCL5, cytotoxic mediators including GZMB, GZMA and TRAIL (TNFSF10), and TNF-superfamily ligands including LIGHT (TNFSF14), TNFSF13B and TNFSF12. Additional interactions involved immune-regulatory and adhesion-associated molecules, including ENTPD1 and ICAM1. These predicted signals were directed towards multiple myeloid, dendritic, epithelial and B-lineage populations, supporting the potential participation of GZMB⁺ CD4⁺ T cells in broad inflammatory, chemotactic, cytotoxic and immunoregulatory circuits within the mucosal microenvironment.

Incoming ligand-receptor interactions that were enriched in GZMB⁺ CD4⁺ T cells were dominated by HLA class I- and class II-associated signals arising from GC and memory B cells, macrophages, CD8⁺ T-cell subsets and epithelial populations (Supplementary Figure 3D). CCL5-mediated interactions from CD8+ intraepithelial T cells were also preferentially enriched. Together, these findings suggest that GZMB⁺ CD4⁺ T cells are likely to occupy a microenvironment characterised by enhanced antigen presentation.

In summary, these findings define a transcriptionally distinct polyfunctional cytotoxic CD4⁺ T-cell state in inflamed UC mucosa. This state is characterised by coordinated cytotoxic, inflammatory and immunoregulatory programmes, acquisition of a late effector phenotype, and responsiveness to cytokine and immunoregulatory cues. GZMB⁺ CD4⁺ T cells also exhibited preferential intercellular communication with lymphocyte, myeloid, and epithelial populations, consistent with their likely integration within an antigen-rich inflammatory microenvironment and a potential role in propagating mucosal immune responses.

### 4. Polyfunctional cytotoxic CD4⁺ T cells are spatially organised within the UC colonic mucosa

Given the distinct intercellular communication patterns associated with the CD4^+^ T cells exhibiting a polyfunctional cytotoxic effector state, we next sought to define their spatial organisation within the inflamed UC colonic microenvironment. To address this, we performed single-cell resolution spatial transcriptomics on colonic samples from 16 patients with active UC (Supplementary Table 2).

Using label transfer of CD4^+^ T-cell populations from the scIBD resource, we identified seven CD4⁺ T-cell subsets in this spatial dataset: activated T cells, terminally differentiated effector memory cells (Temra), T follicular helper (Tfh) cells, Th17 cells, tissue central memory cells (tissue-Tcm), naïve T cells (Tn) and tissue-resident memory cells (Trm) (Supplementary Figures 4A and 4B).

**Figure 4:**
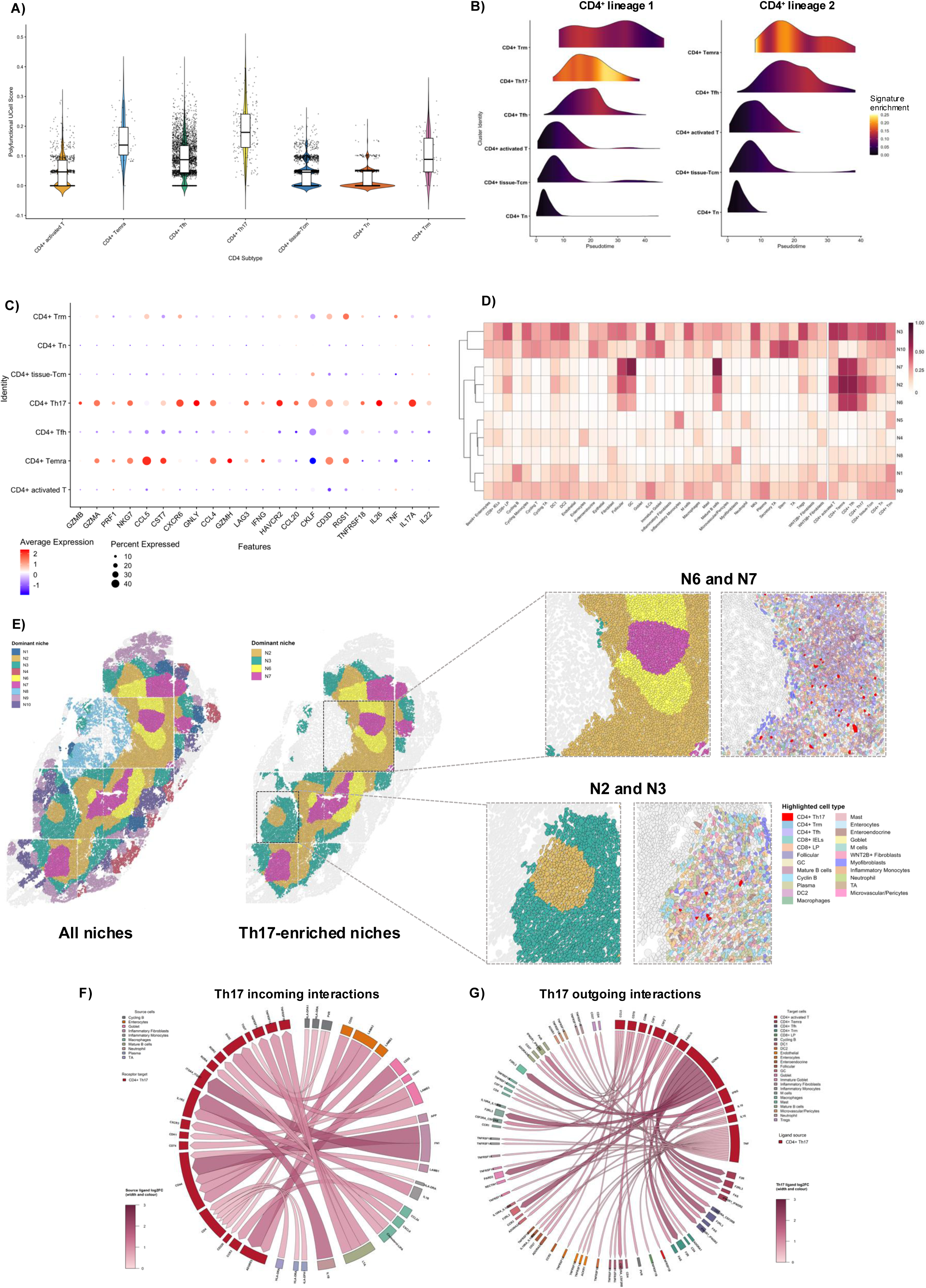
The polyfunctional cytotoxic CD4⁺ T-cell programme is spatially organised across lymphoid and inflammatory niches in UC mucosa. **A,** Single-cell enrichment of the polyfunctional cytotoxic CD4⁺ T-cell gene signature across spatially resolved CD4⁺ T-cell subsets from 16 patients with active UC, quantified using UCell. Each point represents one cell. Each subset was compared with the pooled remaining CD4⁺ populations using linear mixed-effects models incorporating participant identity as a random intercept, with Benjamini-Hochberg correction across comparisons. **B,** Distribution of CD4⁺ T-cell subsets along two Slingshot-inferred differentiation trajectories rooted in naïve CD4⁺ T cells. Ridge height represents cell density across pseudotime and colour denotes polyfunctional cytotoxic gene signature enrichment. **C,** Expression of genes constituting the polyfunctional cytotoxic CD4⁺ T-cell gene signature represented in the CosMx panel along with TNF, IL17A and IL22, across spatially resolved CD4⁺ T-cell subsets. Dot size represents the percentage of cells expressing each gene and colour represents scaled average expression. **D,** Relative distribution of annotated cell populations across the 10 CD4⁺ T-cell-centred spatial niches identified by k-means clustering of cell-type proportions within 50-µm neighbourhoods. For each cell type (column), values represent the proportion of all cells of that type assigned to each niche and were min–max scaled across niches. Darker shading therefore indicates niches containing a greater relative proportion of the corresponding cell type within each niche. Niches were hierarchically clustered for visualisation. **E,** Spatial organisation of the identified niches in a representative UC colonic tissue section. The complete niche map is shown on the left, followed by selective visualisation of the Th17-enriched Niches 2, 3, 6 and 7. Enlarged regions show the the N7 lymphoid core surrounded by the N6 perifollicular region and the broader N2-N3 immune-inflammatory microenvironments. Corresponding cell maps highlight CD4⁺ Th17 cells in red and neighbouring lymphoid, myeloid, epithelial, stromal and vascular populations according to the indicated colours. **F,** Spatially constrained CellChat analysis showing incoming ligand–receptor interactions predicted to be stronger in CD4⁺ Th17 cells than in CD4⁺ Tissue Tcm cells. Source cell populations and their ligands are connected to cognate receptors expressed by CD4⁺ Th17 cells. **G,** Outgoing ligand-receptor interactions predicted to be stronger from CD4⁺ Th17 cells than from CD4⁺ Tissue Tcm cells, connecting Th17-derived ligands to cognate receptors on lymphoid, myeloid, stromal, vascular and epithelial populations. In **F** and **G**, only interactions significant in the Th17 network, with a positive communication-probability difference and differential ligand expression, are shown. Ribbon width and colour intensity represent the ligand log₂ fold change, and sectors are coloured by source- or target-cell identity GC, germinal centre; IEL, intraepithelial lymphocyte; LP, lamina propria; Tcm, central memory T cell; Temra, terminally differentiated effector memory T cell; Tfh, follicular helper T cell; Tn, naïve T cell; Trm, tissue-resident memory T cell; UC, ulcerative colitis.

To determine which spatially resolved CD4⁺ T-cell populations expressed the cytotoxic programme identified in the preceding analysis, enrichment of the polyfunctional cytotoxic CD4⁺ T-cell signature derived from scIBD was quantified at single-cell resolution using UCell (Figure 4A). After accounting for repeated measurements within participants, Th17 cells demonstrated the strongest relative enrichment compared with the mean of the remaining CD4⁺ T-cell populations (estimated difference in enrichment = 0.149, 95% CI 0.121-0.177, adjusted p < 0.0001; Supplementary Table 3). Temra cells were also significantly enriched (estimated difference = 0.095, 95% CI 0.058-0.132, BH-adjusted p < 0.0001). In contrast, activated T cells, tissue-Tcm cells and naïve T cells exhibited significantly lower signature enrichment than the remaining CD4⁺ populations. Collectively, these findings localise the polyfunctional cytotoxic programme to Th17 cells, with additional enrichment observed within Temra cells.

Pseudotime analysis, rooted in CD4^+^ Tn cells, was subsequently performed revealing two inferred differentiation trajectories (Figure 4B and Supplementary Figure 4C). Along Lineage 1, enrichment of the polyfunctional cytotoxic signature increased most prominently within Th17 cells, whereas along Lineage 2, the highest enrichment was observed within Temra cells. Tfh cells showed heterogeneous enrichment across both trajectories, while activated T cells, tissue-Tcm cells and naïve T cells generally exhibited lower scores. This suggests that acquisition of the polyfunctional cytotoxic programme accompanies progression towards differentiated effector states, particularly within Th17 and Temra populations.

Expression of the individual signature genes was subsequently examined alongside the cytokines found to be preferentially enriched in cytotoxic CD4^+^ T cells in our flow cytometry analysis (Figure 4C). Th17 cells demonstrated coordinated expression of cytotoxic effector genes, including *GZMB, GZMA, PRF1, NKG7 and GNLY*, together with inflammatory cytokines and chemokines including *IFNG, TNF, IL17A, IL22, IL26, CCL4, CCL5* and *CCL20,* and immune checkpoint or regulatory molecules, including *HAVCR2, LAG3* and *TNFRSF18*. Although Temra cells also expressed a prominent cytotoxic programme, the concurrent expression of cytotoxic effectors and cytokines was most evident among Th17 cells, further indicating that Th17 cells are most likely to harbour a polyfunctional cytotoxic state within inflamed UC colonic tissues.

CD4⁺ T-cell-centred spatial neighbourhood analysis identified 10 distinct cellular niches (Figure 4D). CD4⁺ Th17 cells were most strongly represented within Niches 2, 3, 6 and 7, which comprised compositionally distinct but spatially connected lymphoid and inflammatory microenvironments (Supplementary Figure 4E). Niche 7 showed the strongest relative enrichment for follicular, germinal centre and mature B cells together with Tfh cells, consistent with a lymphoid core of tertiary lymphoid structures (TLS) (Figure 4D). Niche 6 retained enrichment of B-cell lineage and Tfh populations and spatially surrounded Niche 7, indicative of a perifollicular region surrounding the TLS. Niche 2 contained a broader mixture of CD4⁺ and CD8⁺ T-cell subsets, B-cell lineage cells and antigen presenting populations, forming an immune-rich interface around these lymphoid structures. Niche 3 comprised a more heterogeneous multicellular inflammatory compartment containing lymphoid, myeloid, epithelial, and stromal populations. Spatial mapping demonstrated a nested organisation in which Niche 7 regions were surrounded by Niche 6 and bordered by the broader Niche 2 and Niche 3 compartments (Figure 4E). Th17 cells exhibited some of the highest polyfunctional cytotoxic signature scores across these microenvironments, whereas enrichment within Temra, Tfh and Trm populations was more variable (Supplementary Figure 4E).

Spatial ligand-receptor analysis was then performed to identify candidate signalling interactions involving Th17 cells compared to other CD4 T cell types. In particular, we compared Th17 cells with CD4+ Tcm and CD4+ Tn cells which demonstrated the least enrichment of the polyfunctional cytotoxic signature. Relative to both reference populations, Th17 cells displayed stronger predicted incoming interactions from lymphoid, myeloid, stromal and epithelial populations (Figure 4F and Supplementary Figures 5A and 6A). These included LTA-, CXCL13- and HLA class II-associated interactions involving follicular, germinal centre and mature B-cell populations. Myeloid and tissue-derived signals included chemokines such as CCL19, CXCL9, CCL24 and CCL28, inflammatory/immunoregulatory ligands including IL-1β, MIF and LGALS9, and extracellular matrix or adhesion-associated ligands including FN1 and NECTIN2. These predicted incoming signals are consistent with the localisation of Th17 cells across both TLS-associated N6/N7 compartments and the broader inflammatory N2/N3 interface.

Th17 cells also exhibited increased predicted outgoing communication across multiple effector pathways relative to both Tcm and Tn cells (Figures 4G and Supplementary Figures 5B and 6B). These included chemokines such as CCL5, CXCL13, CCL20 and CCL3, inflammatory cytokines and TNF-superfamily ligands including IFN-γ, IL-16, TNF, LTA, OSM and LIGHT, and cytotoxic mediators including GZMA, FASLG and TRAIL. These interactions were directed towards T- and B-cell, myeloid, stromal, vascular and epithelial populations. Collectively, the spatial ligand-receptor analysis positions polyfunctional cytotoxic Th17 cells at the interface between local antigen-rich lymphoid microenvironments and broader multicellular inflammatory niches, supporting a central role for these cells in orchestrating mucosal inflammation in ulcerative colitis.

### 5. The polyfunctional cytotoxic CD4⁺ T-cell state associates with mucosal disease severity and biologic treatment resistance in UC patients

Given the broad effector properties of this T-cell state, we next examined its clinical relevance by assessing its association with disease severity and therapeutic outcomes in UC patients. The polyfunctional cytotoxic CD4⁺ T-cell signature was evaluated across independent UC cohorts using gene set variation analysis (GSVA)^14^.

In the Mount Sinai Crohn’s and Colitis Registry^15^, this signature was significantly enriched in rectal biopsies from UC compared with non-IBD controls (N = 300 vs 225; p = 7.9e-12, Mann-Whitney U test) (Figure 5A) and increased progressively with endoscopic disease severity (Figure 5B). We next examined whether this signature was associated with broader differences in the mucosal transcriptome. Principal component analysis (PCA) demonstrated that signature enrichment was also associated with the major axes of mucosal transcriptomic variation among UC samples (Figure 5C). Samples with high and low signature enrichment occupied partially distinct regions of the PCA space, with the strongest separation evident along PC1. After adjustment for endoscopic severity, age and sex, signature enrichment remained independently associated with 23.5% of overall transcriptomic variation (PERMANOVA, permutation p < 0.001). Importantly, this association persisted after excluding genes comprising the signature, with signature enrichment accounting for 10.2% of variation among the remaining highly variable genes (PERMANOVA, permutation p < 0.001). These findings indicate that the polyfunctional cytotoxic CD4⁺ T-cell signature captures a major molecular axis of mucosal heterogeneity in UC that is not attributable solely to variation in its constituent genes and is not fully accounted for by endoscopic disease severity, age or sex.

**Figure 5:**
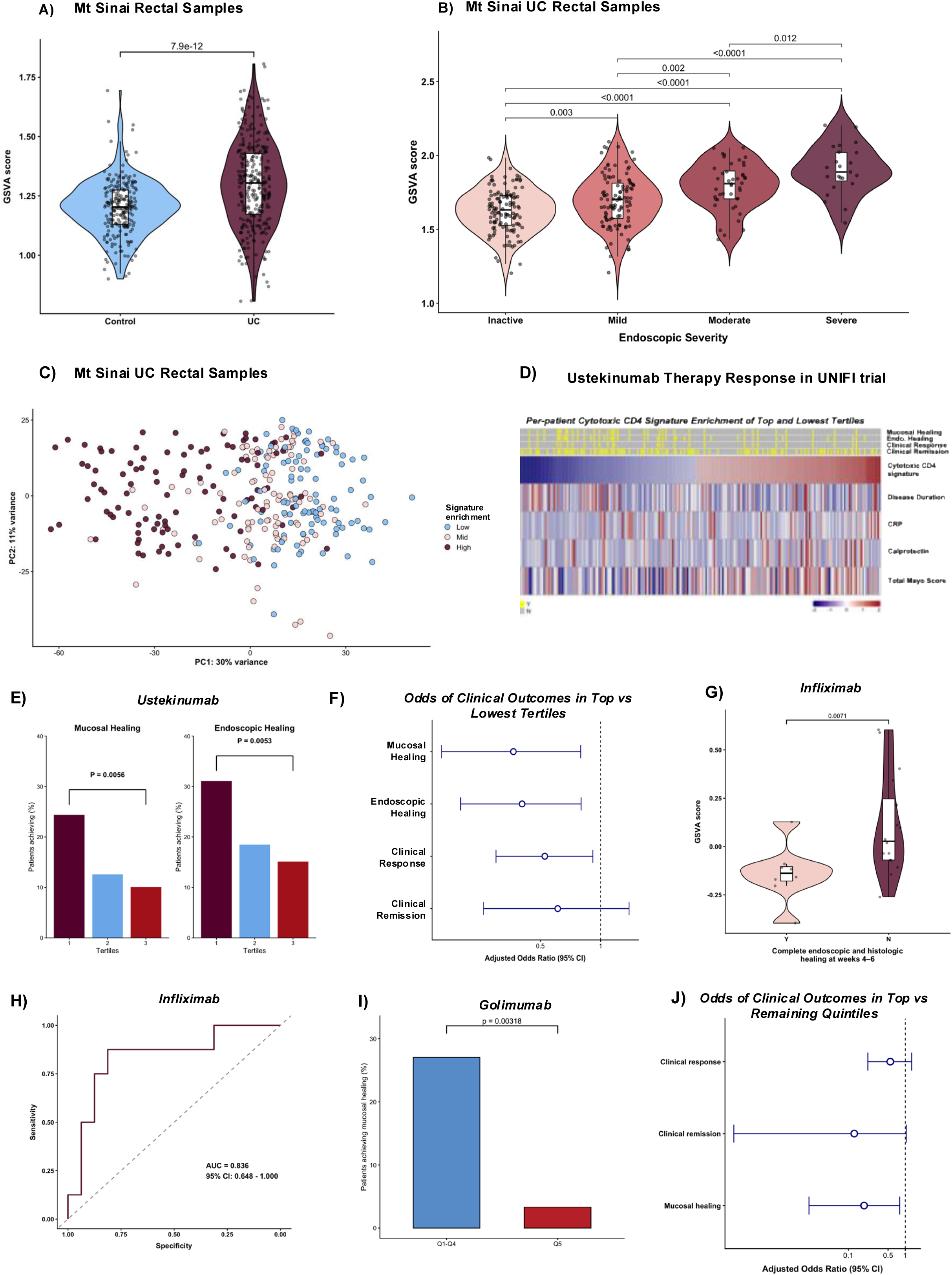
The polyfunctional cytotoxic CD4⁺ T-cell signature is associated with mucosal disease severity and attenuated treatment response to biologic therapies in UC. **A,** Polyfunctional cytotoxic CD4⁺ T-cell signature enrichment, quantified by gene set variation analysis (GSVA), in rectal biopsies from patients with UC (n = 300) and non-IBD controls (n = 225) in the Mount Sinai Crohn’s and Colitis Registry. Groups were compared using a two-sided Mann-Whitney U test. **B,** Signature enrichment across inactive, mild, moderate and severe endoscopic disease categories within the Mount Sinai UC cohort. Pairwise comparisons were performed using two-sided Mann–Whitney U tests with Benjamini-Hochberg correction; adjusted *P* values are shown. **C,** Principal-component analysis of the 500 most variable genes in Mount Sinai UC rectal biopsies. Each point represents one sample and is coloured according to low, intermediate or high tertiles of signature enrichment. **D,** Per-participant heatmap showing treatment outcomes, polyfunctional cytotoxic CD4⁺ T-cell signature enrichment and baseline clinical or inflammatory characteristics among ustekinumab-treated participants in the highest and lowest signature tertiles of the UNIFI trial. Binary outcomes are shown in yellow for achievement and grey for non-achievement of clinical outcome. **E,** Proportions of participants achieving week 8 mucosal healing or endoscopic healing across increasing tertiles of baseline signature enrichment. The lowest and highest tertiles were compared using two-sided Fisher’s exact tests. **F,** Adjusted odds ratios and 95% confidence intervals for week 8 outcomes in the highest relative to the lowest signature tertile. Estimates were obtained using Firth penalised logistic regression adjusted for prior anti-TNF exposure, baseline C-reactive protein, total Mayo score and disease duration. The dashed line indicates an odds ratio of 1. **G,** Baseline signature enrichment in infliximab-treated patients who did or did not achieve complete endoscopic and histological healing at weeks 4-6 in the GSE16879 cohort (n = 24). Groups were compared using a two-sided Mann-Whitney U test. **H,** Receiver-operating-characteristic (ROC) curve evaluating discrimination of complete endoscopic and histological healing using baseline signature enrichment in the infliximab cohort. The area under the curve and 95% confidence interval are shown. **I,** Proportion of golimumab-treated participants in the PROgECT trial achieving week 6 mucosal healing in the highest signature quintile compared with the lower four quintiles combined. Groups were compared using a two-sided Fisher’s exact test. **J,** Adjusted odds ratios and 95% confidence intervals for week 6 clinical response, clinical remission and mucosal healing in the highest signature quintile relative to the lower four quintiles. Estimates were obtained using Firth penalised logistic regression adjusted for baseline total Mayo score and disease duration. The dashed line indicates an odds ratio of 1. AUC, area under the curve; CRP, C-reactive protein; GSVA, gene set variation analysis; OR, odds ratio; UC, ulcerative colitis.

**Figure 6:**
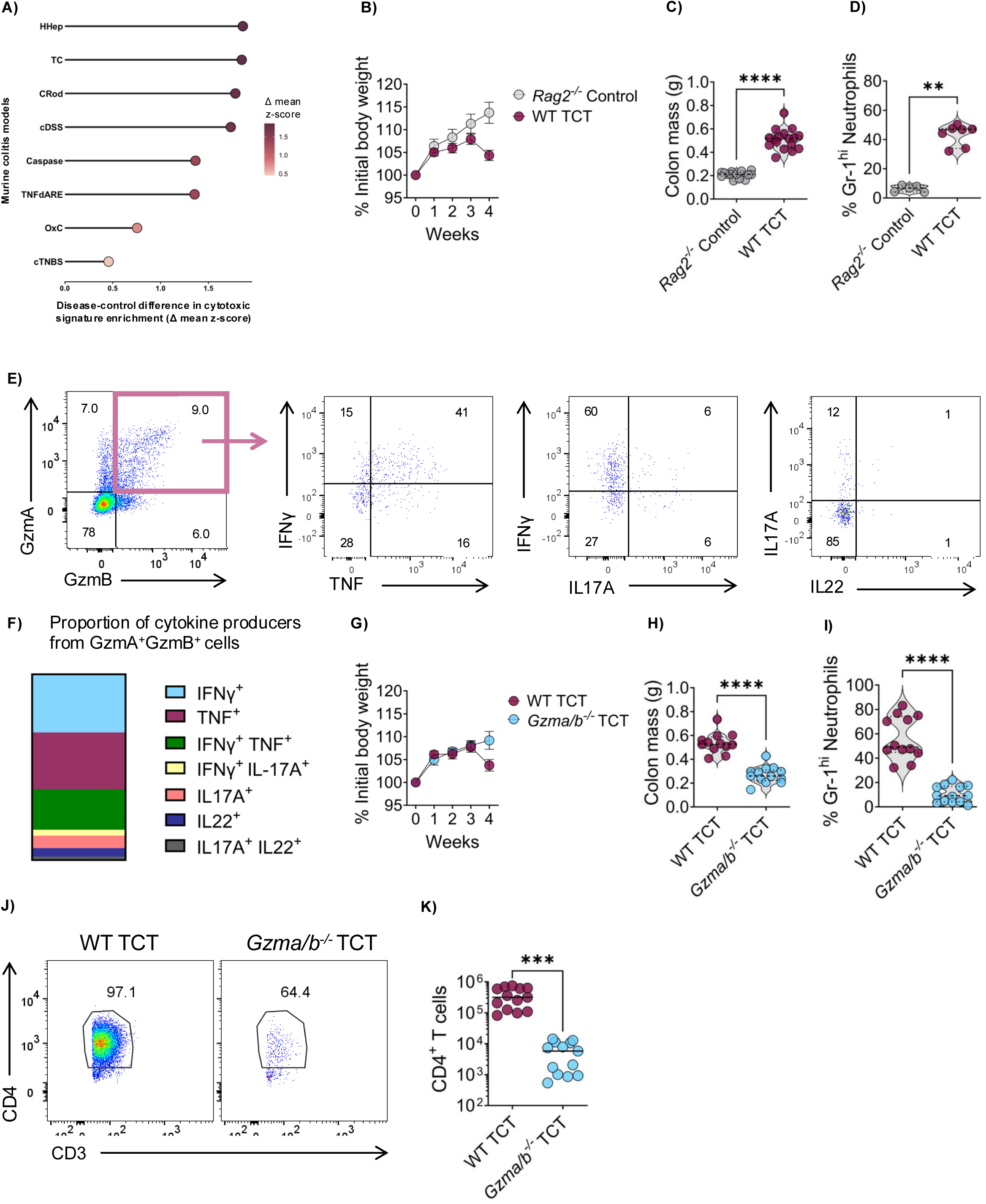
GZMA/GZMB-dependent effector activity is required for development of CD4⁺ T-cell-mediated colitis. **A,** Relative enrichment of the murine orthologues of the polyfunctional cytotoxic CD4⁺ T-cell signature across eight experimental colitis models in the IBD mouse model biobank. Signature enrichment was quantified by GSVA, z-standardised within each model and expressed as the difference between mean disease and control scores. This cross-model comparison was descriptive. **B–D,** Development of colitis following transfer of WT naïve CD4⁺ T cells into *Rag2*⁻/⁻ recipients. **B,** Body weight relative to baseline during the four-week experiment. Data show mean ± s.e.m. **C,** Colon mass and **D,** frequency of colonic Gr-1^hi neutrophils at the experimental endpoint. *Rag2*⁻/⁻ mice not receiving T cells served as controls. **E,** Representative flow cytometry identification of GZMA⁺GZMB⁺ donor-derived CD4⁺ T cells recovered from recipient mice and their expression of IFN-γ, TNF, IL-17A and IL-22. Numbers indicate the percentage of cells within the corresponding gates or quadrants. **F,** Relative composition of mutually exclusive cytokine expression profiles among GZMA⁺GZMB⁺ donor-derived CD4⁺ T cells following WT T-cell transfer. **G–K,** Comparison of colitis following transfer of WT or *Gzma/Gzmb*-deficient naïve CD4⁺ T cells into *Rag2*⁻/⁻ recipients. **G,** Body weight relative to baseline during the four-week experiment. Data show mean ± s.e.m. **H,** Colon mass and **I,** frequency of colonic Gr-1^hi neutrophils at the experimental endpoint. **J,** Representative flow cytometry plots showing donor-derived CD3⁺CD4⁺ T cells recovered from the colon following WT or *Gzma/Gzmb*-deficient T-cell transfer. Numbers indicate the percentage of cells within the indicated gate. **K,** Absolute numbers of donor-derived colonic CD4⁺ T cells. In **C**, **D**, **H**, **I** and **K**, each point represents one mouse and groups were compared using two-sided Mann-Whitney U tests. \*\**P* < 0.01; \*\*\**P* < 0.001; \*\*\*\**P* < 0.0001. cDSS, chronic dextran sodium sulphate colitis; cTNBS, chronic trinitrobenzene sulphonic acid colitis; CRod, *Citrobacter rodentium* infection; HHep, *Helicobacter hepaticus* infection; OxC, oxazolone colitis; TC, CD4⁺ T-cell transfer colitis; TCT, T-cell transfer; TNFΔARE, TNFΔARE colitis; WT, wild type.

We next assessed signature association with response to biologic therapies across independent UC clinical cohorts. Using transcriptomics data^16^ from the UNIFI phase III clinical trial of ustekinumab (anti-IL12/23p40)^17^ (n = 364), patients in the highest tertile of signature enrichment had significantly lower rates of mucosal healing (10% vs 25%, p = 0.0056) and endoscopic healing (14% vs 32%, p = 0.0053) (Figures 5D and 5E). In multivariable analyses adjusted for prior anti-TNF exposure, baseline CRP, total Mayo score and disease duration, high signature enrichment was associated with reduced odds of week 8 mucosal healing (adjusted odds ratio (OR) = 0.37, 95% CI 0.16-0.80, p = 0.011), clinical response (adjusted OR = 0.53, 95% CI 0.30-0.91, p = 0.023) and endoscopic healing (adjusted OR = 0.41, 95% CI 0.20-0.80, p = 0.009).

In the GSE16879 infliximab-treated UC cohort^18^, the signature was significantly enriched in baseline biopsies from patients who failed to achieve complete endoscopic and histologic healing at week 4-6 (p = 0.007; Mann-Whitney U test) (Figure 5G). Baseline signature enrichment discriminated between patients who did and did not achieve complete endoscopic and histological healing, with an AUC of 0.836 (95% CI 0.648-1.000) (Figure 5H). Similarly, in the PROgECT phase II trial of golimumab^19^, week 6 mucosal healing was achieved in only 1 of 30 patient samples (3.3%) in the highest quintile of cytotoxic CD4⁺ T-cell signature enrichment, compared with 33 of 122 samples (27.0%) in the lower four quintiles combined (p = 0.0032; Fisher’s exact test). In multivariable analyses adjusted for baseline total Mayo score and disease duration, the highest enrichment quintile remained associated with significantly lower odds of mucosal healing (adjusted OR = 0.19, 95% CI 0.02-0.80, p = 0.020). A similar association was observed for clinical remission, although this was at the threshold of statistical significance after adjustment (adjusted OR = 0.13, 95% CI 0.001-1.04, p = 0.055). The association with clinical response was attenuated after adjustment (adjusted OR = 0.55, 95% CI 0.22-1.29, p = 0.170; Figure 5J).

Collectively, these findings indicate that enrichment of this transcriptional programme is a marker of reduced response to anti-TNF and anti-IL-12/23p40 therapy, independent of baseline disease severity.

### 6. Genetic disruption of cytotoxic CD4^+^ effector programmes attenuates experimental colitis

We next sought to determine we next sought to determine whether this effector state in CD4+ T cells is functionally required for colitis development *in vivo*. We first surveyed the IBD mouse model biobank^20^, a compendium of transcriptomic datasets derived from multiple murine colitis models under standardised experimental conditions, to identify the *in vivo* models that most closely recapitulate the cytotoxic CD4^+^ T-cell programme observed in human UC. This signature was most enriched in the CD4^+^ T-cell transfer colitis model, as well as in infection-driven colitis models induced by *Helicobacter hepaticus* and *Citrobacter rodentium* (Figure 6A). Subsequently, we used the CD4⁺ T-cell transfer colitis model to investigate the functional contribution of cytotoxic CD4⁺ T-cell effector programmes to colitis (Supplementary Figure 7A). In this model, naïve CD4⁺ T cells are transferred into immunodeficient Rag2⁻/⁻ recipient mice, where their expansion and differentiation induce intestinal inflammation^21^. Expectedly, Rag*2^−/-^* recipients of wild type (WT) CD4⁺ T cells developed severe colitis, characterised by marked weight loss, increased colon mass, splenomegaly and significant increase in neutrophil infiltration (Figures 6B-D and Supplementary Figures 7B and 7C).

**Figure 7:**
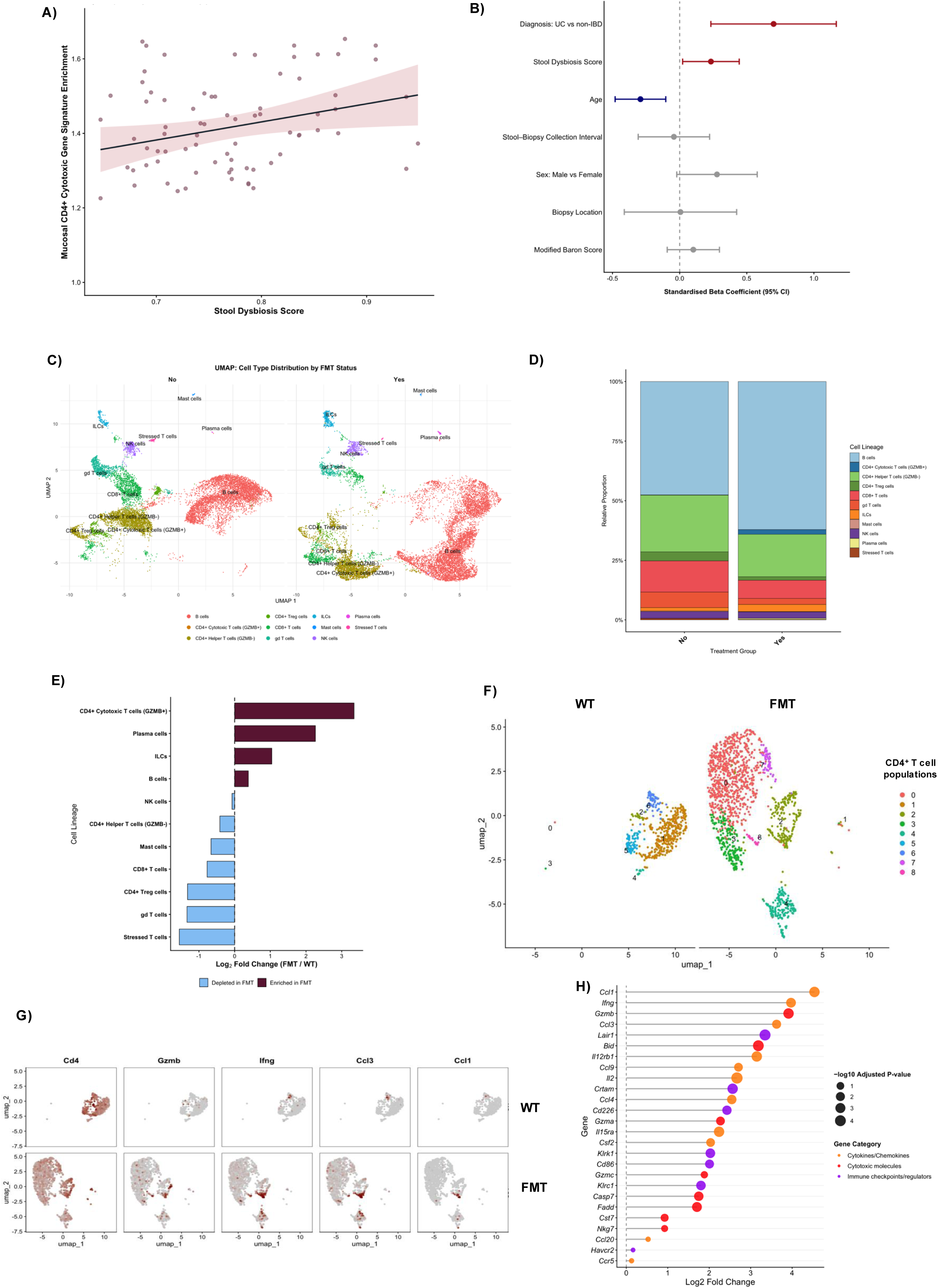
Intestinal dysbiosis is associated with and promotes the emergence of a polyfunctional cytotoxic CD4⁺ T-cell programme. **A,** Association between stool dysbiosis score and mucosal polyfunctional cytotoxic CD4⁺ T-cell signature enrichment in paired host-transcriptomic and stool-metagenomic samples from the IBD Multi-omics Database. The analysis included 77 paired observations from 46 participants. The line shows the fitted unadjusted linear-regression model and shading represents the 95% confidence interval. **B,** Standardised regression coefficients and 95% confidence intervals from a multivariable model of mucosal signature enrichment incorporating diagnosis, stool dysbiosis score, age, stool-biopsy collection interval, sex, biopsy location and Modified Baron endoscopic score. Coefficient estimates used participant-clustered CR2 variance estimation with Satterthwaite-adjusted degrees of freedom. The dashed line indicates no association. **C,** UMAP visualisation of colonic lamina propria immune-cell populations from control and FMT-treated WT mice six weeks after transfer of dysbiotic microbiota from TRUC mice. Cells are coloured according to annotated population. **D,** Descriptive relative composition of the colonic immune-cell compartment in control and FMT-treated mice. **E,** Descriptive log₂ fold changes in cell-population proportions following FMT relative to controls. Positive and negative values indicate relative enrichment and depletion following FMT, respectively. **F,** UMAP visualisation of re-clustered CD4⁺ T cells from control and FMT-treated mice, coloured by transcriptionally defined subpopulation. **G,** Expression of *Cd4, Gzmb, Ifng, Ccl3* and *Ccl1* across the CD4⁺ T-cell UMAPs from control and FMT-treated mice. **H,** Pseudobulk differential-expression analysis of the total CD4⁺ T-cell compartment following FMT relative to controls. The horizontal axis shows log₂ fold change, point size denotes −log₁₀(adjusted *P* value) and colour indicates functional gene category. Displayed genes had log₂ fold change >1 and Benjamini–Hochberg-adjusted *P* < 0.05. FMT, faecal microbiota transfer; IBDMDB, IBD Multi-omics Database; ILC, innate lymphoid cell; TRUC, T-bet⁻/⁻*Rag2*⁻/⁻ ulcerative colitis model; UMAP, uniform manifold approximation and projection; UC, ulcerative colitis; WT, wild type.

Flow cytometry analysis of donor-derived CD4⁺ T cells recovered from recipient mice demonstrated enrichment of GZMA and GZMB, including a population co-expressing both granzymes (Figure 6E and Supplementary Figures 7D and 7E) associated with heterogeneous inflammatory cytokine expression (Figure 6F). The largest components comprised cells expressing IFN-γ or TNF individually, together with a substantial IFN-γ⁺TNF⁺ population. Smaller populations expressed the Th17-associated cytokines IL-17A and IL-22, including IFN-γ⁺IL-17A⁺ and IL-17A⁺IL-22⁺ phenotypes. Thus, double-granzyme-expressing CD4⁺ T cells combined GZMA and GZMB expression with diverse Th1- and Th17-associated cytokine states, recapitulating key features of the polyfunctional cytotoxic CD4⁺ T-cell programme identified in human UC.

We next compared colitis development following adoptive transfer of naïve CD4^+^ T cells isolated from wild-type (WT) BALB/c mice versus *Gzmb/a^−/-^* BALB/c mice, in which granzyme-mediated cytotoxic effector function is genetically ablated, into *Rag2^−/-^* recipient mice (Supplementary Figure 8A). Strikingly, adoptive transfer of *Gzmb/a*^−/-^ deficient CD4⁺ T cells abrogated disease, characterised by absence of weight loss and increased colonic mass alongside significantly reduced neutrophil infiltration and donor CD4⁺ T cells accumulation within the colon (Figures 6G-K and Supplementary Figures 8B and 8C). Importantly, there was absent expression of the polyfunctional cytokine co-expression profile observed following WT cell transfer.

These findings demonstrate a functional requirement for granzyme-dependent CD4⁺ T-cell effector activity in the development of colonic inflammation, implicating the polyfunctional cytotoxic CD4⁺ T-cell programme as a pathogenic mediator of colitis *in vivo*.

### 7. Dysbiotic microbiota promotes the emergence of mucosal polyfunctional cytotoxic CD4⁺ T-cell programmes in the colon

Multimodal evaluation of UC colonic tissues revealed that the polyfunctional cytotoxic CD4+ T-cell state likely arises in the context of heightened antigen-driven stimulation. This was supported by enrichment of HLA-DRB5-associated upstream signals in NicheNet analysis, ligand-receptor interactions implicating heightened MHC class I and II signals acting on cytotoxic CD4⁺ T cells, and robust induction of this programme in infection-driven colitis models. Together, these observations raise the possibility that gut microbial antigen-dependent cues contribute to the emergence of this state. To evaluate this further, we analysed paired host transcriptomic and stool metagenomic data from the IBD Multi-omics Database (IBDMDB)^22^. The analysis included 46 participants, comprising 25 with UC and 21 non-IBD controls, who contributed 49 and 28 colonic mucosal biopsy samples, respectively. Each mucosal transcriptomic sample was paired with the temporally closest stool metagenomic sample collected within four weeks, yielding 77 paired tissue-stool observations. Stool dysbiosis scores calculated by the original IBDMDB study from the metagenomic data was used for this analysis.

Stool dysbiosis score was positively associated with mucosal polyfunctional cytotoxic CD4^+^ T-cell signature enrichment (standardised β = 0.256, 95% CI 0.003-0.510, p = 0.048) (Figure 7A) which persisted after multivariable analyses (Figure 7B). UC diagnosis demonstrated the largest positive association with signature enrichment relative to non-IBD controls (adjusted β = 0.700, 95% CI 0.233-1.167, p = 0.005). After adjustment for diagnosis, stool dysbiosis was the only other covariate demonstrating a statistically significant positive association with the signature (standardised β = 0.233, 95% CI 0.022-0.444, p = 0.033). Increasing age was independently associated with lower signature enrichment (standardised β = -0.292, 95% CI -0.481 to -0.103, p = 0.005). Leave-one-participant-out analysis produced consistently positive dysbiosis coefficients across all iterations, with a median standardised β of 0.234 and a range of 0.164-0.276, indicating that the primary association was not driven by an individual participant. In a sensitivity analysis restricted to UC patients (n=25), the association remained positive and was greater in magnitude, although it no longer reached statistical significance (standardised β = 0.308, 95% CI −0.047 to 0.663, p = 0.082) (Supplementary Table 4).

To determine if dysbiotic microbial communities can induce this phenotype, faecal microbiota transfer (FMT) experiments were performed using colitogenic microbiota from the T-bet^−/-^ RAG2^−/-^ ulcerative colitis (TRUC) model^23^ into WT recipient mice. Single-cell transcriptomic profiling of colonic lamina propria mononuclear cells following FMT demonstrated a marked relative expansion of *Gzmb*⁺ *Cd4*⁺ T cells, which showed the greatest fold increase among immune populations compared with control mice (Figures 7D and 7E). Relative shifts in other immune cell types were also observed, including increases in plasma cells, B cells, and ILCs, alongside proportional reductions in classical cytotoxic populations such as *Cd8*⁺ T cells and NK cells.

Within the *Cd4*⁺ T-cell compartment, a distinct subpopulation co-expressing *Gzmb*, *Ifng* and chemokines including *Ccl3* and *Ccl1* emerged following dysbiotic FMT and was largely absent in control mice (Figures 7F and 7G). Consistent with the emergence and expansion of this population, pseudobulk analysis of the total CD4⁺ T-cell compartment following FMT demonstrated increased expression of cytotoxic effector genes (*Gzmb, Gzma, Gzmc, Casp7, Fadd, Nkg7*), inflammatory cytokines (*Ifng, Il2*), chemokines (*Ccl1, Ccl3, Ccl4, Ccl9*) and immune-regulatory molecules (*Lair1, Crtam, Cd226, Klrk1*) (Figure 7H). This transcriptional shift closely resembled the polyfunctional cytotoxic CD4⁺ T-cell programme observed in human UC colonic tissues. Cytokine receptors including *Il12rb1* and *Il15ra* were also upregulated, consistent with the cytokines identified in human tissue analyses as candidate upstream regulators of this programme.

In summary, these findings link intestinal dysbiosis to the emergence of a polyfunctional cytotoxic CD4⁺ T-cell state within antigen- and cytokine-rich inflammatory niches. This programme was associated with greater mucosal disease severity and reduced biologic treatment response in UC patients, while genetic disruption of granzyme-dependent CD4⁺ T-cell effector activity attenuated colitis *in vivo*.

## DISCUSSION

In this study, we define a distinct microbiota-responsive, spatially organised polyfunctional cytotoxic CD4⁺ T-cell state that promotes mucosal inflammation in UC. By integrating *ex vivo* T-cell-receptor stimulation, multiparameter flow cytometry, bulk, single-cell and spatial transcriptomics, independent clinical cohorts and functional *in vivo* experiments, we identify a CD4⁺ T-cell programme characterised by coordinated cytotoxic, cytokine, chemokine, and immunoregulatory activity. This was preferentially enriched in inflamed UC colonic mucosa, increased with endoscopic disease severity, and was associated with treatment resistance. Crucially, adoptive transfer of *Gzma/Gzmb*-deficient CD4⁺ T cells markedly attenuated experimental colitis and abrogated the polyfunctional cytokine phenotype, implicating granzyme-dependent cytotoxicity as a key effector mechanism of CD4⁺ T-cell-mediated intestinal inflammation.

These findings extend an evolving understanding of cytotoxicity as a pathogenic CD4⁺ T-cell effector programme. Once considered largely a consequence of prolonged *in vitro* activation, cytotoxic CD4⁺ T cells are increasingly recognised in settings of persistent immune stimulation, including chronic infection, malignancy, transplantation and autoimmune disease^24^. More recently, granzyme-expressing CD4⁺ T-cell populations have been described in Crohn’s disease^25^ and murine colitis^26^. Our findings build on these observations by demonstrating that, in UC, cytotoxicity in CD4^+^ T cells forms part of a broader polyfunctional effector state that is spatially organised within the inflamed mucosa, responsive to the intestinal microbiota, and functionally implicated in colonic inflammation.

We observed that this polyfunctional cytotoxic effector state was most enriched within Th17 cells, although components of this programme were also detected in Temra cells. Pseudotime analysis positioned the acquisition of this polyfunctional cytotoxic effector state towards later stages of CD4^+^ T-cell differentiation. Increased expression of *PRDM1*, *RUNX3* and *HOPX*, together with inferred NF-κB-and STAT-associated regulatory activity, was consistent with acquisition of a differentiated effector phenotype. Spatial analysis revealed Th17 cells occupied interconnected microenvironments spanning TLS-like lymphoid cores and perifollicular regions, together with broader multicellular inflammatory niches. Predicted incoming HLA class II-, LTA- and chemokine-associated signals from B-cell lineage and myeloid cell populations were consistent with antigen-rich lymphoid stimulation, whereas outgoing cytotoxic, cytokine and chemokine interactions connected Th17 cells with epithelial, myeloid, stromal, and lymphoid compartments. These findings position polyfunctional cytotoxic CD4⁺ Th17 cells as potential inflammatory hubs coupling local adaptive immune activation to tissue-wide effector responses in UC.

Several complementary analyses implicated inflammatory cytokines and microbial cues in the emergence of this programme. NicheNet, pseudotime and gene regulatory analyses converged on IL-12-, IL-15-, IL-23-, IFN-γ-, EBI3- and TNF-associated signalling, together with antigen-presentation and co-stimulatory pathways. These signals are compatible with the polyfunctional phenotype: IL-12 and IFN-γ can promote Th1-associated effector differentiation, IL-23 maintains pathogenic Th17 responses, and IL-15 supports the survival and cytotoxic differentiation of activated lymphocytes^27,28^. Host-microbiome analyses provided further evidence that this state is microbiota responsive. Stool dysbiosis was associated with mucosal signature enrichment after adjustment for diagnosis and available clinical covariates, while transfer of dysbiotic TRUC microbiota into WT mice induced a *Gzmb*⁺ CD4⁺ T-cell population expressing cytotoxic mediators, inflammatory cytokines, chemokines and cytokine receptors resembling those identified in human UC. These findings support a model in which dysbiotic microbial communities provide antigenic and pro-inflammatory cues to promote this state *in vivo*. However, it is not yet known whether this state is driven by specific microbial antigens, metabolites, epithelial injury or secondary cytokine responses.

Notably, this programme was consistently associated with poor biologic response across UC patient cohorts suggesting that it may mark a treatment-refractory mucosal state. Higher pretreatment signature enrichment associated with poorer outcomes following infliximab, golimumab and ustekinumab therapy, including after adjustment for available baseline covariates of disease severity. Given these therapies target different inflammatory pathways, the T-cell state may therefore reflect a broader inflammatory profile that is insufficiently controlled by blockade of a single cytokine axis. Spatial ligand-receptor analysis identified increased predicted outgoing communication from Th17 cells involving TNF-, LTA- and OSM-associated pathways towards inflammatory monocytes, neutrophils, fibroblasts and endothelial cells. This aligns with previous work implicating OSM-dependent stromal-myeloid circuits in treatment-refractory IBD, raising the possibility that polyfunctional cytotoxic CD4⁺ T cells participate in multicellular inflammatory networks that are incompletely suppressed by individual targeted therapies^29,30^. Prospective validation and assessment of incremental performance beyond established clinical markers will be required to determine whether the programme represents a pan-treatment prognostic marker or a treatment-specific predictive biomarker.

The adoptive T-cell transfer experiments established a functional role for GZMA/GZMB-dependent CD4⁺ T-cell activity in colonic inflammation *in vivo* suggesting that granzyme expression in CD4⁺ T-cells is not merely a marker of T-cell activation but contributes directly to colitis pathogenesis. GZMA and GZMB may contribute to intestinal inflammation through direct cytotoxic activity, extracellular matrix remodelling, regulation of T-cell differentiation or survival, or indirect effects on immune cell recruitment^31^. Direct killing assays using intestinal epithelial cells or patient derived colonic organoids, together with selective disruption of individual granzymes and perforin, will be required to further disentangle these possibilities. Other limitations include the incomplete cellular specificity of the polyfunctional cytotoxic signature, the inferential nature of the trajectory and cell-cell communication analyses, and the retrospective nature of biologic treatment response analyses. These limitations are partly mitigated by the convergence of transcriptomic, spatial, protein-level and experimental evidence, but define important areas for future investigation.

The identification of this state has potential implications for molecular stratification and therapeutic development in UC. A clinically deployable assay of this programme could ultimately help identify patients with a highly activated, multicellular mucosal immune state, although translation will require prospective validation and demonstration of added value over clinical, biochemical, and endoscopic markers. Candidate therapeutic approaches could target upstream microbial or cytokine signals identified in this work, disrupt the tissue niches supporting this state, or selectively modulate its granzyme-dependent effector functions. Such strategies would need to avoid general suppression of cytotoxic immunity, given its importance in antiviral and antitumour responses.

In summary, our findings unveil a microbiota-responsive, spatially organised polyfunctional cytotoxic CD4⁺ T-cell state that is associated with disease severity and reduced biologic response in UC. More broadly, this work supports a framework in which pathogenic immunity in UC is organised not solely around canonical T-helper lineages, but around functional states shaped by microbial exposure, inflammatory signals and the local tissue microenvironment.

## METHODS

### Patient recruitment and sample collection

For the human flow cytometry analysis, a total cohort of 49 participants was prospectively recruited. This included 32 patients with UC who had endoscopically active disease (UCEIS ≥2) and 17 non-IBD controls undergoing routine colonoscopy to investigate non-inflammatory gastrointestinal symptoms. Participants were recruited across two hospital networks: initially at King’s College Hospital / Guy’s and St Thomas’ NHS Foundation Trust (n=41), and subsequently at Imperial College Healthcare NHS Trust (n=8). A subset of patients also provided peripheral blood samples (24 UC and 7 non-IBD controls).

During endoscopy, 7-12 mucosal biopsies were collected from inflamed (UC) or non-inflamed (HC) mucosa into sterile PBS. For the initial KCL cohort, the study protocol was approved by the local R&D departments and the National Institute for Health Research - Research Ethics Committee (REC reference: 15/LO/1998), and was adopted by the South London Clinical Research Network (CRN). For the subsequent Imperial cohort, patient recruitment and sample collection were conducted under the THAMES-IBD study ethical approval (Yorkshire & The Humber - Sheffield REC, Ref: 22/YH/0043). The studies were conducted in accordance with the Declaration of Helsinki.

### Isolation of peripheral blood mononuclear cells from human blood

Peripheral blood mononuclear cells (PBMCs) were isolated from peripheral blood by density-gradient centrifugation using Lymphoprep™ (STEMCELL Technologies) according to the manufacturer’s instructions. Viable cells were enumerated by trypan blue exclusion using 0.4% Trypan Blue Solution (Thermo Fisher Scientific). Freshly isolated PBMCs were either used immediately for downstream applications or cryopreserved in foetal bovine serum (FBS) containing 10% dimethyl sulfoxide (DMSO) and stored in liquid nitrogen until use.

### Isolation of lamina propria mononuclear cells from human colonic biopsies

Colonic biopsy specimens obtained during endoscopy were mechanically disrupted using sterile scalpels and enzymatically digested in RPMI 1640 medium (Gibco) supplemented with 10% foetal calf serum (FCS), 1 mg mL⁻¹ collagenase D, and 10 U mL⁻¹ DNase I (Sigma-Aldrich). Digestion was performed for 1 h at 37 °C using a gentle-MACS Dissociator (Miltenyi Biotec). The resulting cell suspension was further dissociated by passage through a 16-gauge needle and filtered through a 70-μm cell strainer. Mononuclear cells were enriched by density-gradient centrifugation over Lymphoprep™ (STEMCELL Technologies) at 1,000 × g for 20 min at room temperature. Lamina propria mononuclear cells (LPMCs) were collected from the interface and used for subsequent flow cytometric analyses.

### Intracellular cytokine staining and flow cytometry

PBMCs and LPMCs were rested overnight in complete RPMI 1640 medium supplemented with recombinant human IL-2 (100 U/ml), penicillin (100 U/ml), streptomycin (100 μg/ml), gentamicin (20 μg/ml), amphotericin B (2.5 μg/ml), metronidazole (1 μg/ml) and 2-mercaptoethanol (2.5 μl/L). Cells were subsequently stained with antibodies against TCR Vα7.2, TCR γδ and TCR Vα24-Jα18 (BioLegend) for 30 min at 4°C to identify MAIT, γδ T-cell and invariant natural killer T (iNKT) cell populations.

Following TCR staining, cells were stimulated with phorbol 12-myristate 13-acetate (PMA; 50 ng/ml) and ionomycin (1 μg/ml) for 3 h at 37°C in the presence of Brefeldin A and Monensin (Thermo Fisher Scientific). Following stimulation, cells were stained with antibodies against CD3, CD4, CD8, CD161, CD56, integrin β7 and a fixable viability dye (BioLegend and BD Biosciences) for 30 min at 4°C. Cells were then fixed using 1X BD Cytofix solution (BD Biosciences) for 1 h at room temperature, washed, and permeabilized using 1X Perm/Wash buffer (BD Biosciences). Intracellular staining was subsequently performed using antibodies against IL-17A, TNF, IL-22 (BioLegend), granzyme B, and IFN-γ (BD Biosciences). Following staining, cells were washed and resuspended in PBS for acquisition.

Samples were acquired on an LSR II flow cytometer (BD Biosciences) and analysed using FlowJo software v10 (BD/Tree Star). Cytokine-producing cells were identified using unstimulated control samples cultured in the presence of Brefeldin A and Monensin alone. Boolean gating was used to define polyfunctional T-cell populations based on all possible cytokine-expression combinations.

Within GZMB⁺ CD4⁺ T cells, 16 mutually exclusive expression patterns defined by IFN-γ, TNF-α, IL-17A and IL-22 were quantified for each participant. Patterns were aggregated according to the number of cytokines expressed and separately to calculate marginal expression of each cytokine irrespective of co-expression patterns. UC and non-IBD groups were compared using two-sided Mann-Whitney U tests, with Benjamini-Hochberg correction applied separately across the four cytokine-number categories, 16 mutually exclusive patterns, and four marginal cytokine comparisons.

### Isolation of lamina propria cells using a cell-foam walkout culture system

For *in vitro* stimulation assays and transcriptional analyses, LPMCs from 4 UC patients were isolated using a cell-foam matrix walkout culture system as previously described^32^. Briefly, Cellfoam matrices (9 mm × 9 mm × 1.5 mm; Cytomatrix Pty Ltd) were autoclaved and coated with rat-tail collagen I (100 μg/ml; BD Biosciences) in PBS for 30 min at 37 °C, followed by two washes with PBS. Colonic biopsies were partially disrupted by gently compressing the epithelial surface into the collagen-coated matrix. Individual matrices were transferred to separate wells of a 24-well tissue culture plate and cultured in 2 mL complete RPMI 1640 medium containing 10% FCS, β-mercaptoethanol, penicillin (100 U/ml), streptomycin (100 μg/ml), metronidazole (1 μg/ml), gentamicin (20 μg/ml), amphotericin B (2.5 μg/ml), recombinant human IL-2 (100 U/ml; Novartis Pharmaceuticals UK) and recombinant human IL-15 (10 ng/ml; BioLegend). Cultures were maintained at 37 °C in a humidified incubator containing 5% CO₂. Immune cells were allowed to migrate from the embedded tissue into the surrounding culture medium. Following the 48-hour walkout period, recovered LPMCs from four patients with UC were divided into paired stimulated and unstimulated conditions. Cells were incubated for 4 hours in the presence or absence of agonistic anti-CD3 antibody (1μg/ml) stimulation and were lysed in Qiazol afterwards (Qiagen, Germany).

### Bulk RNA-sequencing of *ex vivo* stimulated LPMCs

RNA was subsequently extracted (Qiagen RNeasy mini kit) and sequencing libraries were generated using NEBNext® Ultra™ RNA Library Prep Kit for Illumina® (NEB, USA) following manufacturer’s recommendations. Briefly, mRNA was purified from total RNA using poly-T oligo-attached magnetic beads. Fragmentation was carried out using divalent cations under elevated temperature in NEBNext First Strand Synthesis Reaction Buffer (5X). First strand cDNA was synthesized using random hexamer primer and M-MuLV Reverse Transcriptase (RNase H-). Second strand cDNA synthesis was subsequently performed using DNA Polymerase I and RNase H. Remaining overhangs were converted into blunt ends via exonuclease/polymerase activities. After adenylation of 3’ ends of DNA fragments, NEBNext Adaptor with hairpin loop structure were ligated to prepare for hybridization. Library fragments were purified with AMPure XP system (Beckman Coulter, Beverly, USA) and treated with 3 μl USER Enzyme (NEB, USA) at 37°C for 15 min, followed by 5 min at 95 °C. Then PCR was performed with Phusion High-Fidelity DNA polymerase, Universal PCR primers and Index (X) Primer. Library quality was assessed on Agilent Bioanalyzer 2100 and on Nanodrop ND-1000 Spectrophotometer. The library preparations were sequenced on an Illumina HiSeq platform, generating 150 bp paired end reads^33,34^.

### Animal husbandry

BALB/c (Charles River code: 028) mice, *Gzma* x *Gzmb* double knockout (*Gzmb/a*^−/-^) and *Rag2^−/-^* mice (both Jackson Labs) were obtained from commercial sources. TRUC mice have been described previously^23,35,36^. All animals were housed together under specific pathogen-free conditions at Imperial Hammersmith CBS, or Charles River Laboratories. Experiments involving animals were conducted in accredited facilities, complying with the UK Animals (Scientific Procedures) Act 1986 under Home Office Licence Numbers PPL: PP9922202.

### Isolation of murine colonic lamina propria cells

Isolation of murine colonic lymphocytes has previously been described^37^. Mice were euthanized by a Home Office approved Schedule 1 method and organs were excised in an aseptic manner. Briefly epithelium was removed by incubation in HBSS with 5 mM EDTA and 10 mM HEPES at 37°C for 20 min. Tissue was then homogenized and digested in HBSS with 2% FCS and supplemented with 0.5 mg/ml collagenase D, 10 μg/ml DNase I and 1.5 mg/ml Dispase II (all Roche) for 20 min at 37°C. The digested lymphocyte-enriched population was harvested using a 40%–80% Percoll (GE Healthcare) gradient centrifugation for cLPMCs and a 40%–70% for the cell suspension from the intraepithelial layer.

### Cell sorting of murine colonic lamina propria cells

Splenic cells were isolated into a single-cell suspension in complete animal medium with the use of 70 μM mesh filter and general mechanical destruction. The suspension was centrifuged at 860 *g* for 5 min at 4°C, and red blood cells were lysed using a standard red blood lysis buffer (ACK). CD4^+^ T cells from the spleen of mice were purified using LS positive selection MACs and anti-CD4 (L3T4) beads (Miltenyi Biotec). CD4^+^ MACs sorted cells were further purified by FACS using a FACS Aria (BD Biosciences) with a 70 μm nozzle insert and FACS Diva software. CD4^+^ MACs sorted cells were stained for 20 min at 4°C in the dark with the following antibodies: anti-CD4-PerCPCy5.5 (RM4-5), anti-CD25-PE (PC61), anti-CD62L-PECy7 (MEL-14; Thermo Fisher), and anti-CD44-Pacific Blue (IM1.8.1; Thermo Fisher). Single-positive compensation controls and unstained controls were used to set up instrument settings and for gating strategies. Cell purity was verified post-sort (requiring 95% purification) and cell viability was assessed using trypan-blue staining. Sorted cells were collected in complete media (as described above), counted using a hemocytometer, and immediately cultured or transferred in vivo.

### Naïve T-cell transfer model of colitis

Naïve T cells were transferred as described^37^. Spleens were harvested from either WT BALB/c mice or *Gzmb/a*^−/-^ mice and mechanically disrupted as described above. CD4^+^ T cells with naïve markers (CD4^+^ CD25^−^ CD44^low^ CD62L^high^) were sorted using a FACS Aria to a purity of >95%, washed, and resuspended in sterile PBS. *Rag2^−/−^* mice were injected intraperitoneally with 5 × 10^5^ CD4^+^ cells per mouse and humanely culled after 4 weeks. Mice were monitored for their health every week for signs of illness.

### Gene expression quantification

Fastq files from CD3 stimulated and unstimulated LPMCs were processed by Novogene with in-house Perl scripts to discard reads with adaptor contamination, or at least 10% of uncertain bases (N), or at least 50% of nucleotides with a Phred quality score less than 20. The remaining read pairs were then aligned to the human genome (GRCh37/hg19) using TopHat v2.0.12^33^. Counts of the read pairs mapped uniquely and concordantly to each gene were computed with HTSeq v0.6.1^34^.

Fastq files from whole tissue biopsies were processed with trimmomatic (v. 0.39)^38^ to remove adaptor contamination and poor-quality bases were. The resulting read files were mapped to the GRCh38 assembly of the human genome using Hisat2^39^ with default parameters. The number of reads mapping to the genomic features annotated in Ensembl^40^ with a MAPQ score higher than or equal to 10 was calculated for all samples using htseq-count (v. 0.11.3)^34^ with default parameters. Data for Ensembl genes with no associated ENTREZ gene identifier were discarded; the read counts for Ensembl genes mapped to the same ENTREZ gene identifier were summed up sample wise.

### Differential expression and pathway enrichment analyses

Differential gene expression analysis of paired CD3-stimulated and unstimulated LPMCs was performed in R (version 4.5.2) using DESeq2 (version 1.50.2)^41^. Gene-level counts were modelled using a negative-binomial generalised linear model incorporating donor identity and stimulation condition. Genes with fewer than 10 cumulative reads were excluded, and unstimulated samples were specified as the reference. Differential expression was assessed using Wald tests with Benjamini–Hochberg correction. Differentially expressed genes were defined by an absolute log₂ fold change >1 and adjusted *P* <0.05. Variance-stabilised counts were used for principal-component analysis. Ensembl identifiers were mapped to HGNC symbols and Entrez Gene identifiers using clusterProfiler (version 4.18.4)^42^, AnnotationDbi (version 1.72.0) and org.Hs.eg.db (version 3.22.0).

Preranked gene-set enrichment analysis was performed using clusterProfiler with the fgsea backend (version 1.36.2). Genes were ranked by unshrunken log₂ fold change. Where multiple features mapped to the same Entrez identifier, the feature with the greatest absolute fold change was retained. Gene Ontology Biological Process and human KEGG pathways were analysed using gene-set sizes of 5-500 and 10-500 genes, respectively. Pathways with Benjamini-Hochberg-adjusted p-value < 0.05 were considered significant, with enrichment direction determined by the normalised enrichment score.

### IBD TaMMA analysis

The reproducibility of cytotoxic transcriptional programmes was evaluated using the IBD Transcriptome and Metatranscriptome Meta-Analysis (TaMMA) resource^8^, comprising harmonised bulk transcriptomic data from 26 studies of IBD patients. TaMMA differential expression results for UC versus tissue-matched non-IBD controls were downloaded directly from the platform and analysed separately for colonic and rectal tissue in R (version 4.5.2). Ensembl identifiers were mapped to Entrez identifiers using org.Hs.eg.db (version 3.22.0), with log₂ fold changes averaged where multiple Ensembl identifiers mapped to the same Entrez identifier. Genes were ranked by decreasing log₂ fold change, and Gene Ontology Biological Process enrichment was performed using clusterProfiler (version 4.18.4), restricting analysis to gene sets containing 10-500 genes. Prespecified pathways were extracted from the complete enrichment results and considered significant at false-discovery rate (FDR) <0.05. Enrichment direction and magnitude were summarised using normalised enrichment scores.

For gene-level analysis, harmonised TaMMA expression data were obtained for eight prespecified cytotoxic effector genes: *GZMB, GZMA, GZMH, PRF1, GNLY, NKG7, FASLG* and *CRTAM*. Analyses were restricted to UC and non-IBD control samples from the colon and rectum. Expression values were transformed as log₂(counts + 1), and UC and control samples were compared separately within each tissue using two-sided Mann-Whitney U test.

### scIBD single-cell transcriptomic analysis

Publicly available single-cell RNA-sequencing data, cell annotations, clinical metadata and UMAP coordinates were obtained from the scIBD resource. Analyses were restricted to adult colonic, rectal or hindgut samples from inflamed UC samples. Data were analysed in R (version 4.5.2) using Seurat (version 5.3.0)^43^ and SeuratObject (version 5.3.0). The original scIBD UMAP coordinates and cell annotations were retained. CD4⁺ T cells were isolated using the annotated Th17, Tfh, Temra, Trm, activated, tissue-Tcm and naïve T-cell populations.

Cells with normalised log-transformed *GZMB* expression >0 were classified as *GZMB*⁺. Differential expression between *GZMB*⁺ and *GZMB*⁻ CD4⁺ T cells from inflamed UC tissue was performed using MAST (version 1.36.0)^44^ through Seurat’s FindMarkers function, incorporating patient identity as a latent variable and requiring gene detection in ≥5% of either population. The 20 most strongly upregulated genes meeting log₂ fold change >1 and adjusted *P* <0.05 were used to define the polyfunctional cytotoxic CD4⁺ T-cell signature. For pathway analysis, genes were ranked by average log₂ fold change and mapped to Entrez identifiers using org.Hs.eg.db (version 3.22.0). Gene Ontology Biological Process and Reactome enrichment were evaluated using clusterProfiler (version 4.18.4) and ReactomePA (version 1.54.0)^45^, respectively. Pathways with Benjamini-Hochberg-adjusted p <0.05 were considered significant.

Trajectory inference was performed using the largest contributing UC dataset from Smillie *et al.*^12^ and was restricted to annotated CD4⁺ memory, activated Fos-low, activated Fos-high, PD-1⁺ and MT-high populations from donors containing ≥10 *GZMB*⁺ cells. The 2,000 most variable genes were used for scaling, principal-component analysis and UMAP generation using the first 30 principal components. Lineages were inferred using Slingshot (version 2.18.0)^46^, with CD4⁺ memory cells specified as the starting population. Pseudotime distributions were compared between *GZMB*⁺ and *GZMB*⁻ cells using one-sided Wilcoxon rank-sum tests. Gene expression dynamics were modelled along each lineage using LOESS smoothing (span = 0.75), evaluated at 50 equally spaced pseudotime positions and standardised to gene-level z-scores.

Transcription factor activity associated with the GZMB⁺ CD4⁺ T-cell state was inferred using VIPER implemented in decoupleR (version 2.12.0)^47^. The complete vector of MAST-derived average log₂ fold changes was used as the input molecular signature. The human DoRothEA regulatory network (downloaded 06/10/2025)^48^ was restricted to high- and intermediate-confidence interactions (levels A-C), with a minimum regulon size of five target genes. Positive and negative VIPER scores indicated inferred increases and decreases in transcription factor activity, respectively, in GZMB⁺ relative to GZMB⁻ CD4⁺ T cells. Regulators with absolute activity scores greater than 1.5 were prioritised for visualisation.

Candidate upstream ligands were identified using NicheNet^10^ and its 2021 human ligand-receptor and ligand-target networks. Cell populations from inflamed UC tissue were treated as candidate senders and *GZMB*⁺ CD4⁺ T cells as receivers. Ligands were prioritised according to their expression in sender populations, receptor expression in receiver cells, predicted ligand-target regulatory potential, Pearson correlation and area under the precision-recall (AUPR) curve.

Cell-cell communication was additionally inferred in inflamed UC samples using LIANA^13^ with the CellPhoneDB method. *GZMB*⁺ and *GZMB*⁻ CD4⁺ T cells were treated as separate sender and receiver populations. Interactions with CellPhoneDB-derived *P* <0.05 were retained, and communication magnitude was represented by the LIANA lr_means score. For each ligand-receptor pair and partner population, differential communication was calculated as lr_means for *GZMB*⁺ CD4⁺ T cells minus that for *GZMB*⁻ CD4⁺ T cells; interactions absent from one state were assigned a value of zero. Positive values therefore indicated preferential communication involving *GZMB*⁺ CD4⁺ T cells. Incoming and outgoing interactions were summarised using heatmaps and chord diagrams.

### IBD Multi-omics Database analysis

Paired host transcriptomic and stool metagenomic data were obtained from the IBD Multi-omics Database (IBDMDB)^22^. Participants with UC or non-IBD controls who had both data types available were retained. Each mucosal transcriptomic sample was paired with the temporally closest stool metagenomic sample obtained from the same participant within a maximum interval of four weeks. Where multiple stool samples met this criterion, the sample with the smallest absolute collection time difference was selected. The corresponding stool dysbiosis score, calculated by the original IBDMDB study, was used for analysis. Analyses were restricted to colonic and rectal biopsy samples.

Enrichment of a predefined 20-gene GZMB-associated cytotoxic T-cell signature was quantified in each mucosal transcriptomic sample using single-sample gene set enrichment analysis (ssGSEA), implemented using GSVA (version 2.4.7)^14^. The association between stool dysbiosis score and mucosal signature enrichment was first examined using an unadjusted linear regression model. A multivariable linear regression model was subsequently fitted, adjusting for diagnosis, age, sex, colonic biopsy location, Modified Baron endoscopic score and the stool-biopsy collection interval. Modified Baron codes with absent values for non-IBD controls were assigned a value of zero.

Analyses were performed using complete cases, with non-IBD status, female sex and rectal biopsy location specified as the reference categories for categorical variables.

The outcome and continuous predictors were z-standardised, allowing associations to be reported as standardised regression coefficients. To account for repeated observations from individual participants, coefficient estimates were evaluated using participant-clustered CR2 variance estimation with Satterthwaite-adjusted degrees of freedom, implemented using clubSandwich (version 0.7.0). Results are presented with cluster-robust 95% confidence intervals and two-sided p values. Sensitivity analyses included restriction to participants with UC, with continuous variables re-standardised within this subgroup. The stability of the primary dysbiosis association was assessed using leave-one-participant-out analysis, in which each participant was sequentially excluded and the multivariable model refitted.

### Analysis of public bulk UC transcriptomic datasets

The polyfunctional cytotoxic CD4⁺ T-cell signature was evaluated in four independent UC cohorts obtained from the Gene Expression Omnibus (GEO): GSE16879^18^, GSE212849^49^, GSE206285^16^ and GSE193677^15^. Sample-level enrichment scores were calculated using single-sample gene-set enrichment analysis implemented in GSVA (version 2.4.7), using the signature genes represented in each dataset.

GSE16879 comprised colonic mucosal microarray profiles from 24 patients with UC obtained before their first infliximab treatment^18^. Raw CEL files were background-corrected and normalised using robust multi-array averaging with the oligo package. Probe sets with median expression greater than 4 and an unambiguous gene annotation were retained, and multiple probes mapping to the same gene were collapsed to a single gene-level value. Baseline signature scores were compared between patients who did and did not achieve complete endoscopic and histological healing 4–6 weeks after treatment using a two-sided Wilcoxon rank-sum test. Discrimination was evaluated using receiver-operating-characteristic analysis, with the area under the curve and its 95% confidence interval calculated using pROC (version 1.19.0.1).

GSE206285^50^ comprised baseline colonic microarray profiles from the UNIFI phase III trial^17^. Processed robust multi-array average expression profiles and clinical metadata were obtained from GEO, and the analysis was restricted to the 364 participants receiving ustekinumab. Week 8 outcomes included clinical response, clinical remission, endoscopic healing and mucosal healing. Signature scores were divided into tertiles, and outcome frequencies in the highest and lowest tertiles were compared using Fisher’s exact tests. Associations were subsequently evaluated using complete-case Firth penalised logistic-regression models adjusted for prior anti-TNF exposure, baseline C-reactive protein, total Mayo score and disease duration.

GSE212849^49^ comprised baseline colonic-biopsy microarray profiles from golimumab-treated participants in the PROgECT phase II trial^19^. Probe sets were mapped to HGNC gene symbols, with duplicate mappings collapsed by mean expression. Week 6 outcomes included clinical response, clinical remission and mucosal healing, with mucosal healing defined as a Mayo endoscopic subscore of 0 or 1. Signature scores were divided into quintiles, and outcomes in the highest quintile were compared with those in the lower four quintiles using Fisher’s exact tests. Adjusted associations were estimated using complete-case Firth penalised logistic-regression models incorporating baseline total Mayo score and disease duration.

For GSE193677^15^, raw rectal-biopsy RNA-sequencing counts and clinical metadata from the Mount Sinai Crohn’s and Colitis Registry were analysed. The analysis included 300 UC and 225 non-IBD control samples. Genes with fewer than 10 cumulative reads were excluded, and counts were variance-stabilised using DESeq2 (version 1.50.2). Signature enrichment was compared between UC and control samples using a two-sided Wilcoxon rank-sum test. Within UC, pairwise comparisons across endoscopic-severity categories were performed using Wilcoxon rank-sum tests with Benjamini-Hochberg adjustment. Principal-component analysis of the UC samples was performed using the 500 most variable genes. The independent association between continuous signature enrichment and the PC1-PC2 coordinate space was assessed by marginal PERMANOVA using Euclidean distances and 9,999 permutations, adjusting for endoscopic severity, age and sex. As a sensitivity analysis, all signature genes were excluded before selecting the 500 most variable genes, and PERMANOVA was repeated on the resulting expression matrix using the same covariates. Unless otherwise specified, statistical tests were two-sided.

### Analysis of IBD mouse model biobank transcriptomics data

Transcriptomic datasets from eight experimental colitis models were obtained from the IBD mouse model biobank: CD4⁺ T-cell transfer colitis, chronic dextran sodium sulphate (DSS) colitis, chronic trinitrobenzene sulphonic acid (TNBS) colitis, TNFΔARE colitis, intestinal epithelial cell-specific *Casp8* deletion, *Helicobacter hepaticus* infection, *Citrobacter rodentium* infection and oxazolone-induced colitis (ArrayExpress accessions E-MTAB-14306, E-MTAB-14329, E-MTAB-14325, E-MTAB-14318, E-MTAB-14316 and E-MTAB-14312)^51^. For each model, transcriptomic profiles from diseased mice were compared with the corresponding control condition. Sample identifiers were matched to the associated metadata, duplicated metadata entries were removed and count matrices were reordered to ensure concordance with sample annotations.

Within each model, genes with fewer than 10 total counts across the relevant samples were excluded. Count data were processed using DESeq2 (version 1.50.2) and variance-stabilising transformation was performed independently for each disease-control comparison. Murine signature gene symbols were mapped to Ensembl identifiers using clusterProfiler (version 4.18.4) and the org.Mm.eg.db annotation database (version 3.22.0). Ensembl version suffixes were removed, and duplicated Ensembl identifiers were excluded.

Enrichment of the available murine orthologues of the polyfunctional cytotoxic CD4⁺ T-cell signature (*n* = 17) was quantified in individual samples using single-sample gene-set enrichment analysis implemented in GSVA (version 2.4.7). Enrichment scores were z-standardised within each experimental model. Mean standardised scores were calculated separately for diseased and control animals, and relative cytotoxic-signature enrichment was defined as the difference between the mean disease and control scores. Experimental models were ranked according to this disease-control enrichment difference. This cross-model comparison was descriptive and was used to prioritise an experimental system for subsequent functional investigation.

### *In vivo* faecal microbiota transplant (FMT) experiment and single cell RNA sequencing

FMT of mice has previously been described within the lab^52^. In brief, faecal content was extracted from the caecum of TRUC mice and reconstituted in sterile PBS with 25% glycerol, which was then orally gavaged into Balb/c mice (200μl). Mice were euthanised after 6 weeks and colonic lamina propria cells extracted as described earlier. Cells were then stained and live CD45^+^ cells were sorted using a FACS Aria machine (BD Biosciences) and taken immediately to be run on the 10x. Cells were suspended at 1×10^6^/mL in PBS and 10,000 cells were loaded onto the ChromiumTM Controller instrument within 15 min after completion of the cell suspension preparation using GemCode Gel Bead and Chip, all from 10x Genomics (Pleasanton, CA), and following the manufacturer’s recommendations. Briefly, cells were partitioned into Gel Beads in Emulsion in the ChromiumTM Controller instrument where cell lysis and barcoded reverse transcription of RNA occurred. Libraries were prepared using 10x Genomics Library Kits and sequenced on an Illumina NextSeq500 according to the manufacturer’s recommendations. Read depth of more than 200 million reads per library, or an approximate average of 10,000 reads per cell was obtained.

### Single-cell RNA-seq data analysis of *in vivo* FMT experiment

The raw 10X Genomics sequencing libraries were processed using the Cell Ranger^53^ suite v.3.0.1 to demultiplex base call files, generate single cell feature counts for each library, and finally combine these data into one feature by barcode matrix. Read alignment and gene expression quantification made use of the CellRanger pre-built mouse (mm10 v. 3.0.0) reference data. The individual UMI count matrices were normalised to the same effective sequencing depth before they were aggregated.

Downstream analyses were performed in R using Seurat (version 5.3.0). The analysis was restricted to FMT-treated and control mice, and sample metadata were matched using unique sample identifiers. Dimensionality reduction and clustering were performed following sample-level integration using Harmony. A shared-nearest-neighbour graph was constructed using the first 30 Harmony dimensions, followed by clustering at a resolution of 0.5 and UMAP visualisation. Cell populations were annotated using canonical lineage markers and cluster-specific differentially expressed genes.

Cell type proportions were calculated relative to the total number of retained cells in each experimental group. Conventional CD4⁺ T cells were identified by detectable *Cd3e* and *Cd4* expression and absence of detectable *Cd8a* expression. Within this population, cells with normalised *Gzmb* expression greater than zero were classified as *Gzmb*⁺ CD4⁺ T cells. Descriptive changes in cellular composition were calculated as the log₂ ratio of population proportions in FMT-treated relative to control mice, using a pseudocount of 0.001.

For focused analysis, CD4⁺ T cells were reprocessed using the 2,000 most variable genes, followed by scaling, principal-component analysis, neighbour identification and UMAP using the first 15 principal components. Raw counts were subsequently aggregated by individual mouse to generate pseudobulk CD4⁺ T-cell expression profiles. Counts were TMM-normalised using edgeR (version 4.8.2), transformed using voom and compared between FMT-treated and control mice using limma (version 3.66.0) with empirical-Bayes moderation. *P* values were adjusted using the Benjamini-Hochberg procedure, and genes with a log₂ fold change greater than 1 and an adjusted p value < 0.05 were considered significantly upregulated following FMT.

### Single-cell resolution spatial transcriptomics

#### Sample preparation

CosMx Spatial Molecular Imaging (SMI) was performed on formalin-fixed, paraffin-embedded (FFPE) colonic biopsies from patients with ulcerative colitis (UC, n=16). FFPE samples were acquired under the THAMES-IBD study ethical framework (Yorkshire & The Humber - Sheffield REC, Ref: 22/YH/0043). FFPE specimens were initially assessed by H&E staining to verify tissue preservation and pathological representation. Samples meeting quality thresholds for both RNA integrity and histomorphological assessment were selected for downstream spatial gene expression analysis.

Tissue sections (5 μm) were mounted onto Leica Bond Plus slides (Leica Biosystems) and baked overnight at 60°C to enhance tissue adhesion. Sections were deparaffinized using xylene and graded ethanol washes, followed by target retrieval at 100°C for 15 minutes. Tissue permeabilization was achieved using Proteinase K (3 μg/mL; NanoString Technologies) for 30 minutes at 40°C.

Following permeabilisation, slides were washed in 2× SSC-T and DEPC-treated water before incubation with fiducial markers for image registration. Samples were subsequently post-fixed in 10% neutral buffered formalin and treated with Sulfo-NHS-Acetate to reduce non-specific binding. After additional washes in 2× SSC, *in situ* hybridization was performed overnight (16–18 hours) at 37°C using the CosMx Human Universal Cell Characterization Panel (NanoString Technologies), which comprises approximately 6,000 RNA targets.

Following stringent washes with SSC and 100% formamide, nuclei were stained with DAPI alongside the cell segmentation markers PanCK, CD45, CD3, and CD298/B2M (Bruker Corporation). Slides were washed in PBS and stored in 2× SSC until imaging. Image acquisition was carried out using the CosMx SMI platform with manufacturer-recommended pre-bleaching and cell-segmentation settings.

#### Primary Processing, Quality Control, and Dataset Subsetting

CosMx Spatial Molecular Imager (SMI) data were processed using the NanoString–Bruker CosMx/AtoMx analysis pipeline. Cell segmentation was performed using Cellpose-based algorithms, with detected transcripts assigned to individual segmented cells. Following primary processing, cell-level expression matrices, cellular metadata, and spatial coordinate information were exported for downstream analysis in R (version 4.3.2).

Downstream analysis was performed using Seurat (version 5.3.0). Quality control (QC) was conducted at both cellular and field-of-view (FOV) levels. Cell-level QC metrics included the proportion of background negative control probes, cell area, and the number of detected genes and transcripts per cell. FOV-level quality assessment included evaluation of fluidic alignment and transcript assignment efficiency. FOVs demonstrating poor transcript localisation, defined as a mean unassigned transcript fraction >20%, were excluded from subsequent analysis.

#### Dimensionality reduction, cell annotation and functional scoring

Expression data were normalised using SCTransform v2 and integrated across samples before UMAP dimensionality reduction using a cosine-distance metric. CD4⁺ T-cell annotations derived from the scIBD resource were transferred to the integrated dataset, identifying activated, Temra, Tfh, Th17, tissue-Tcm, naïve and Trm populations. Assignments were validated using canonical lineage and activation markers.

Enrichment of the 19 genes from the polyfunctional cytotoxic CD4⁺ T-cell signature represented in the spatial panel was quantified in individual cells using UCell (version 2.6.2). Each CD4⁺ subset was compared with the pooled remaining CD4⁺ populations using a linear mixed-effects model incorporating participant identity as a random intercept. P values were adjusted across subset comparisons using the Benjamini-Hochberg procedure.

#### Pseudotime analysis

CD4⁺ T-cell differentiation trajectories were inferred using Slingshot (version 2.18.0), with refined cell-state annotations supplied as cluster labels and naïve CD4⁺ T cells specified as the starting population. Lineage-specific changes in signature enrichment and gene expression were modelled across pseudotime using generalised additive models with cubic regression splines. For visualisation, pseudotime was divided into 50 equal-width bins, and mean scores were calculated for each CD4⁺ subset in bins containing at least five cells. Cell-density distributions and signature dynamics were visualised using normalised ridge plots and LOESS smoothing (span = 0.5), respectively.

#### Spatial neighbourhood analysis

Spatial neighbourhoods were constructed separately within each tissue specimen using cellular centroid coordinates converted to physical distances with the CosMx scaling factor of 0.1202809 µm/pixel. Cells within 50 µm of each focal CD4⁺ T cell were identified using RANN (version 2.6.2); focal cells with fewer than five neighbours were excluded.

Neighbourhood composition was defined by the proportions of annotated neighbouring cell types. Recurring cellular microenvironments were identified by k-means clustering of the neighbourhood-composition matrix (*k* = 10). Mean cell-type proportions were calculated for each niche and min-max scaled for visualisation.

#### Spatial ligand-receptor analysis

Spatially constrained ligand-receptor interactions were inferred using CellChat^54^. Normalised RNA expression, curated cell-type annotations and global cellular coordinates were supplied to CellChat. For the focused UC analysis, cells within 50 μm of a CD4⁺ Th17 or CD4⁺ Tn cell were identified using RANN, with searches performed independently within each slide and FOV. Pixel distances were converted using a scaling factor of 0.12028 μm per pixel. Cell types represented by fewer than 10 cells were excluded, while non-focal populations were randomly downsampled to a maximum of 3,000 cells. All focal CD4⁺ T cells were retained.

Spatial CellChat objects were constructed using the human CellChatDB ligand-receptor database^54^. Overexpressed signalling genes and interactions were identified before communication probabilities were calculated using a 10% truncated mean. Communication was constrained to an interaction range of 50 μm, with contact-dependent interactions restricted to 10 μm. Significance was assessed using 100 bootstrap iterations and interactions involving cell populations with fewer than 10 cells were removed before pathway-level probabilities and aggregated communication networks were calculated.

Outgoing signalling was defined with CD4⁺ Th17 cells as the ligand-producing population, whereas incoming signalling was defined with CD4⁺ Th17 cells as the receptor-expressing population. Th17 communication probabilities were compared separately with CD4⁺ Tn and tissue-Tcm cells by matching cell partner, ligand, receptor and signalling pathway. Differential communication was defined as the Th17 probability minus the corresponding reference-cell probability. Only interactions significant in the Th17 network (*p* ≤ 0.05) and with a positive probability difference were retained. Autocrine interactions, direct interactions between the focal and reference CD4⁺ populations, low-confidence cell annotations, and collagen interactions were excluded.

Ligand expression was evaluated using cell-level differential-expression analysis with Seurat’s Wilcoxon rank-sum test, using a minimum detection fraction of 0.05 and no initial fold-change threshold. For outgoing signalling, CD4⁺ Th17 cells were compared with the corresponding reference population across the complete UC dataset. For incoming signalling, each source-cell population was compared between cells located within 50μm of Th17 cells and cells located within 50 μm of the reference population. These analyses were performed separately by source-cell type, with cells falling within both neighbourhoods excluded and at least 10 cells required in each group. Ligands were considered supported when the log₂ fold change was ≥0.25 and the adjusted *p* value was ≤0.05. Multisubunit ligands were evaluated conservatively using the minimum subunit log₂ fold change and maximum adjusted p value.

Bubble heatmaps displayed receiving or source-cell populations as rows and ligands as columns. Circle area represented the number of unique receptors associated with each ligand, while colour represented ligand log₂ fold change on a continuous scale from 0 to 3. Circos plots displayed retained ligand–receptor relationships, with ribbon thickness and colour intensity scaled continuously according to ligand differential-expression log₂ fold change.

#### Spatial visualisation

Tissue-wide cellular maps were generated from CosMx cell-segmentation boundaries using sf and ggplot2. Polygon identifiers were harmonised with Seurat cell identifiers. Duplicate or incomplete vertices were removed, and cells represented by fewer than three valid vertices were excluded. Boundary coordinates were grouped by cell, closed to form polygon rings, converted to sf geometries and validated before annotation with the corresponding cell type, niche assignment and expression metadata. Regions of interest were selected by specifying the slide and constituent FOVs, with global coordinates used to preserve their relative tissue positions.

Cellular polygons were displayed using geom_sf. Non-highlighted cells were shown in grey, while selected cell populations were assigned fixed colours that were maintained across figures. Individual cellular boundaries were delineated using thin black outlines. Niche-membership maps were produced using the same geometries, with cells coloured according to their dominant niche assignment and unassigned cells displayed in grey. Where individual niches were highlighted, outlines were applied to all constituent cellular polygons rather than only the CD4⁺ niche-centre cells.

Niche-composition heatmaps were generated using pheatmap. Cell counts were normalised within each cell type by dividing the number assigned to each niche by the total number of that cell type across the complete UC dataset. Where applied, proportions were log₂-transformed and rescaled between 0 and 1 for visualisation. Niches were hierarchically clustered, while cell-type ordering was retained across figures to support direct comparison. Niche-proximity networks were constructed using igraph, with nodes representing CD4⁺-centred niches. Node size reflected the number of niche centres, node colour followed the fixed niche palette and edge width represented the normalised fraction of neighbouring niche centres.

Ligand-receptor bubble heatmaps were rendered using ComplexHeatmap, circlize and grid. Rows represented interacting source or receiving cell types and columns represented ligands. Circle area was proportional to the number of unique receptors associated with each ligand, while fill colour represented the corresponding ligand differential-expression log₂ fold change on a continuous scale. The analytical criteria used to select ligand–receptor interactions are described in the CellChat methods section.

Circos diagrams were generated using circlize. Individual ligand and receptor molecules were represented as sectors, grouped and coloured according to their associated cell population. Ribbons connected cognate ligand–receptor pairs, with ribbon width, sector size and continuous colour intensity scaled according to ligand differential-expression log₂ fold change. Extreme values were capped at the 95th percentile to prevent a small number of interactions dominating the visual scale.

### Statistical analysis

Unless otherwise stated, statistical analyses were performed in R (version 4.5.2). Participants and individual mice were treated as biological replicates for human and animal experiments, respectively. Independent groups were compared using two-sided Mann-Whitney U tests. Paired analyses were performed as specified in the corresponding methodological subsections. Where multiple related hypotheses were tested, p values were adjusted using the Benjamini-Hochberg procedure. Multivariable analyses were performed using complete cases and are reported with 95% confidence intervals. Statistical significance was defined as p or adjusted p <0.05, as applicable.

## Supplementary Figure Legends

**Supplementary Figure 1:**
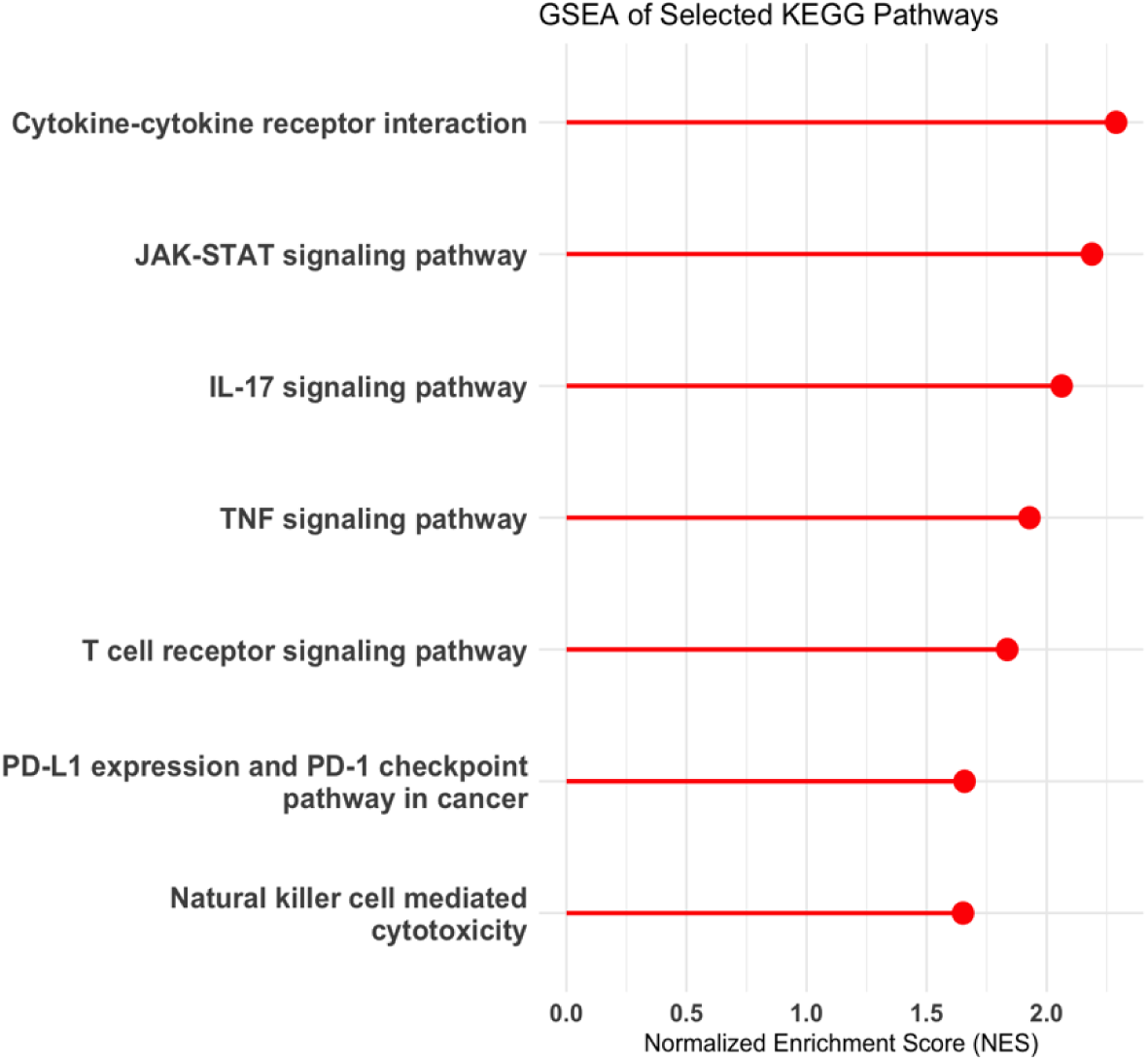
T-cell receptor activation induces inflammatory and cytotoxic KEGG pathways in UC LPMCs. Gene set enrichment analysis of selected Kyoto Encyclopedia of Genes and Genomes (KEGG) pathways following anti-CD3 stimulation of patient-derived colonic LPMCs relative to unstimulated cells. Lines and points indicate the normalised enrichment score (NES); positive values represent enrichment following anti-CD3 stimulation. All displayed pathways met a Benjamini-Hochberg-adjusted *P* value threshold of <0.05. GSEA, gene set enrichment analysis; LPMC, lamina propria mononuclear cell; NES, normalised enrichment score; UC, ulcerative colitis.

**Supplementary Figure 2:**
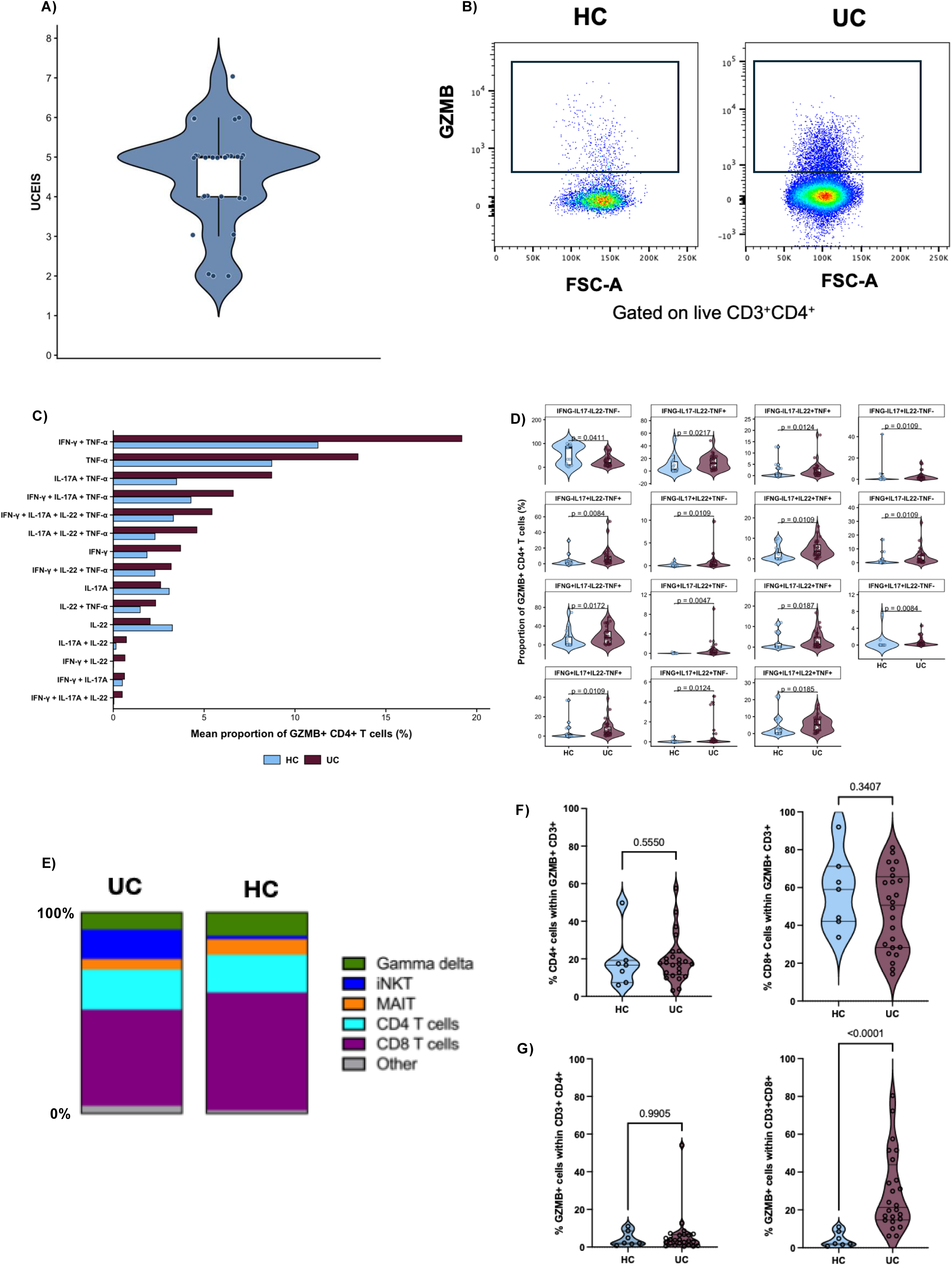
Flow cytometry analysis of mucosal and circulating GZMB-expressing T-cell populations in UC and non-IBD controls. **A,** Distribution of Ulcerative Colitis Endoscopic Index of Severity scores among patients with active UC included in the mucosal flow cytometry analysis (n = 32). Each point represents one participant; the embedded box plot denotes the median and interquartile range. **B,** Representative flow cytometric identification of GZMB⁺ cells among live mucosal CD3⁺CD4⁺ T cells from a non-IBD control and a patient with UC. **C,** Mean proportions of mucosal GZMB⁺ CD4⁺ T cells exhibiting each of the 15 cytokine-positive expression profiles defined by IFN-γ, TNF, IL-17A and IL-22. **D,** Participant-level comparison of the corresponding mutually exclusive cytokine-expression profiles between UC and non-IBD tissues. Each point represents one participant; violin plots show the distributions and embedded box plots denote the median and interquartile range. Groups were compared using two-sided Mann-Whitney U tests, with Benjamini–Hochberg correction across all 16 mutually exclusive profiles, including the cytokine-negative population. Adjusted *P* values are shown. **E,** Mean relative composition of the circulating GZMB⁺CD3⁺ T-cell compartment in UC patients and non-IBD controls. **F,** Proportions of circulating GZMB⁺CD3⁺ T cells belonging to the CD4⁺ and CD8⁺ lineages. **G,** Frequencies of GZMB⁺ cells within circulating CD3⁺CD4⁺ and CD3⁺CD8⁺ T-cell populations. Circulating analyses included participants with available PBMC samples (UC, n = 24; non-IBD controls, n = 7). In **F** and **G**, each point represents one participant and groups were compared using two-sided Mann-Whitney U tests; exact *P* values are shown. HC, non-IBD control; iNKT, invariant natural killer T cell; MAIT, mucosal-associated invariant T cell; PBMC, peripheral blood mononuclear cell; UC, ulcerative colitis; UCEIS, Ulcerative Colitis Endoscopic Index of Severity.

**Supplementary Figure 3:**
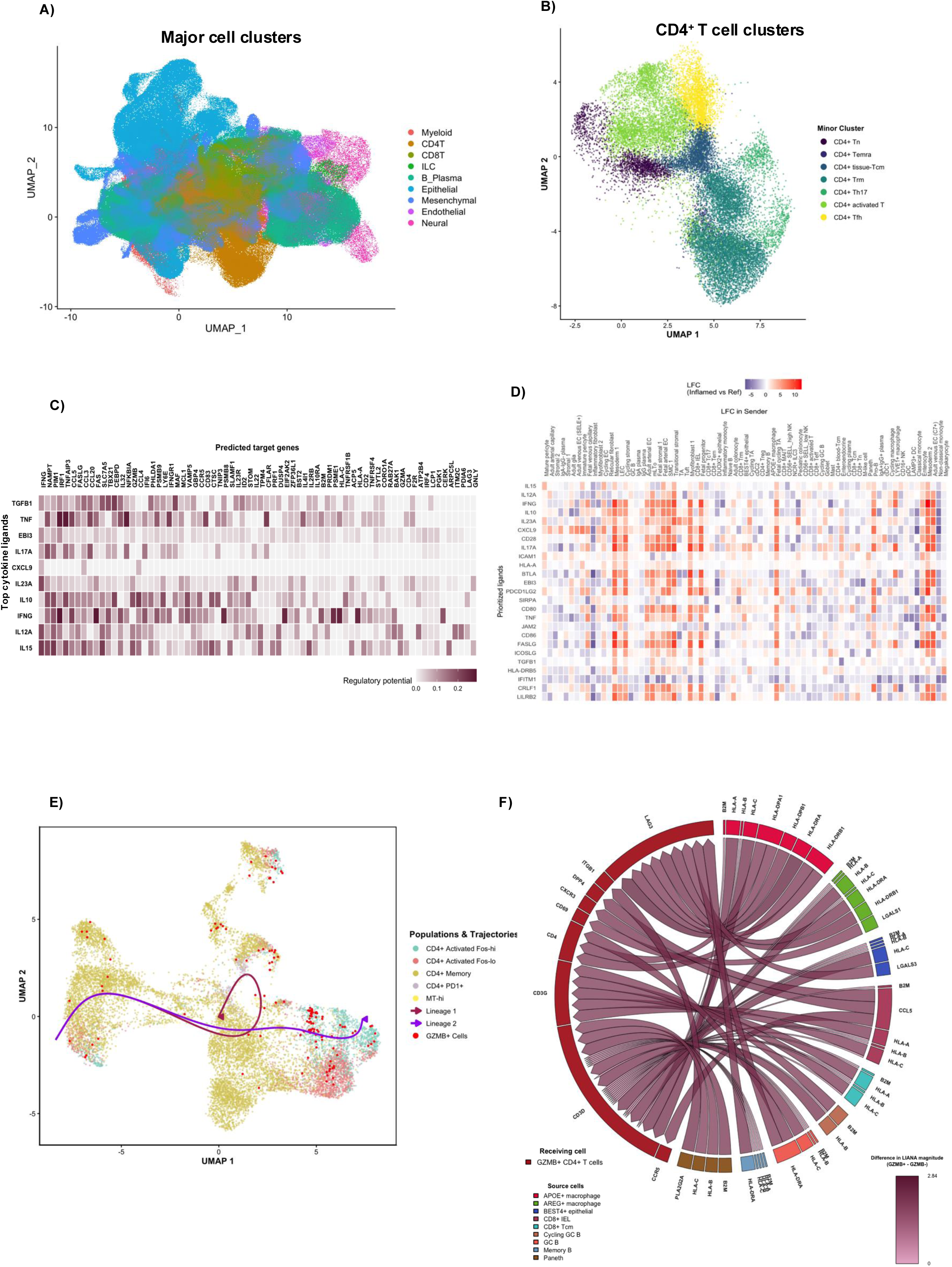
Single-cell characterisation of candidate regulatory signals, differentiation trajectories and incoming communication associated with GZMB⁺ CD4⁺ T cells. **A,** UMAP visualisation of major cellular compartments in inflamed UC colonic tissues from the scIBD resource, coloured according to the original major cell type annotations. **B,** UMAP visualisation of the seven annotated CD4⁺ T-cell populations retained for downstream analysis. **C,** NicheNet ligand-target regulatory matrix showing the predicted regulatory potential of prioritised cytokine ligands for genes comprising the GZMB⁺ CD4⁺ T-cell transcriptional programme. Greater colour intensity indicates higher predicted regulatory potential. **D,** Expression of prioritised NicheNet ligands across candidate sender-cell populations. Colour represents the log₂ fold change in ligand expression in inflamed UC relative to reference tissue within each sender population. **E,** UMAP visualisation of the Smillie *et al.* CD4⁺ T-cell populations used for trajectory inference. Cells are coloured by annotated population, GZMB⁺ cells are highlighted in red and the two Slingshot-inferred trajectories originating from CD4⁺ memory cells are overlaid. **F,** Chord diagram showing selected preferential incoming ligand-receptor interactions received by GZMB⁺ CD4⁺ T cells relative to GZMB⁻ CD4⁺ T cells. Ribbons connect ligands expressed by the indicated source populations with their predicted receptors on GZMB⁺ CD4⁺ T cells; ribbon shading represents the increase in LIANA interaction magnitude relative to GZMB⁻ CD4⁺ T cells. Displayed interactions met a CellPhoneDB-derived *P* value threshold of <0.05. AUPR, area under the precision-recall curve; GC, germinal centre; IEL, intraepithelial lymphocyte; LFC, log₂ fold change; Tcm, central memory T cell; Temra, terminally differentiated effector memory T cell; Tfh, follicular helper T cell; Tn, naïve T cell; Trm, tissue-resident memory T cell; UC, ulcerative colitis; UMAP, uniform manifold approximation and projection.

**Supplementary Figure 4:**
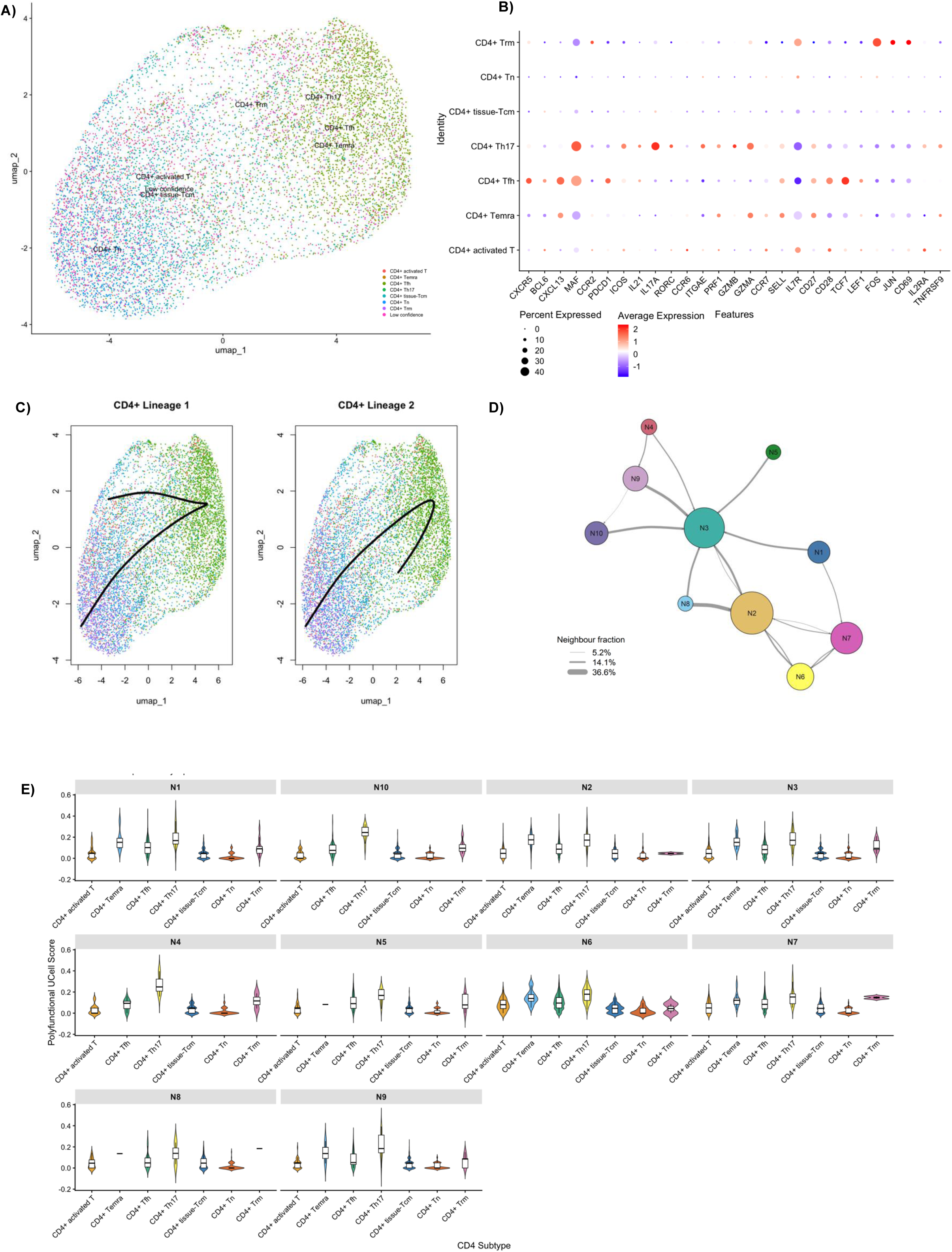
Annotation, trajectory and niche-specific enrichment of spatially resolved CD4⁺ T-cell populations in UC mucosa. **A,** UMAP visualisation of spatially resolved CD4⁺ T cells from 16 patients with active UC following label transfer of CD4⁺ T-cell annotations from the scIBD reference. Cells are coloured according to transferred subtype; cells that could not be assigned confidently are labelled low confidence. **B,** Expression of canonical lineage, differentiation and activation markers across the spatially resolved CD4⁺ T-cell populations. Dot size represents the percentage of cells expressing each gene and colour represents scaled average expression. **C,** Slingshot-inferred CD4⁺ T-cell differentiation trajectories, rooted in naïve CD4⁺ T cells, projected onto the CD4⁺ T-cell UMAP. The two inferred lineages are shown separately. **D,** Spatial adjacency network of the 10 CD4⁺ T-cell-centred niches. Nodes represent individual niche identities and connecting edges indicate spatial neighbourhood relationships; edge width represents the proportion of neighbouring cells assigned to the corresponding niche pair; node size represents relative size of niche. **E,** Cell-level polyfunctional cytotoxic-signature enrichment across CD4⁺ T-cell subsets within each spatial niche, quantified using UCell. Tcm, central memory T cell; Temra, terminally differentiated effector memory T cell; Tfh, follicular helper T cell; Tn, naïve T cell; Trm, tissue-resident memory T cell; UC, ulcerative colitis; UMAP, uniform manifold approximation and projection.

**Supplementary Figure 5:**
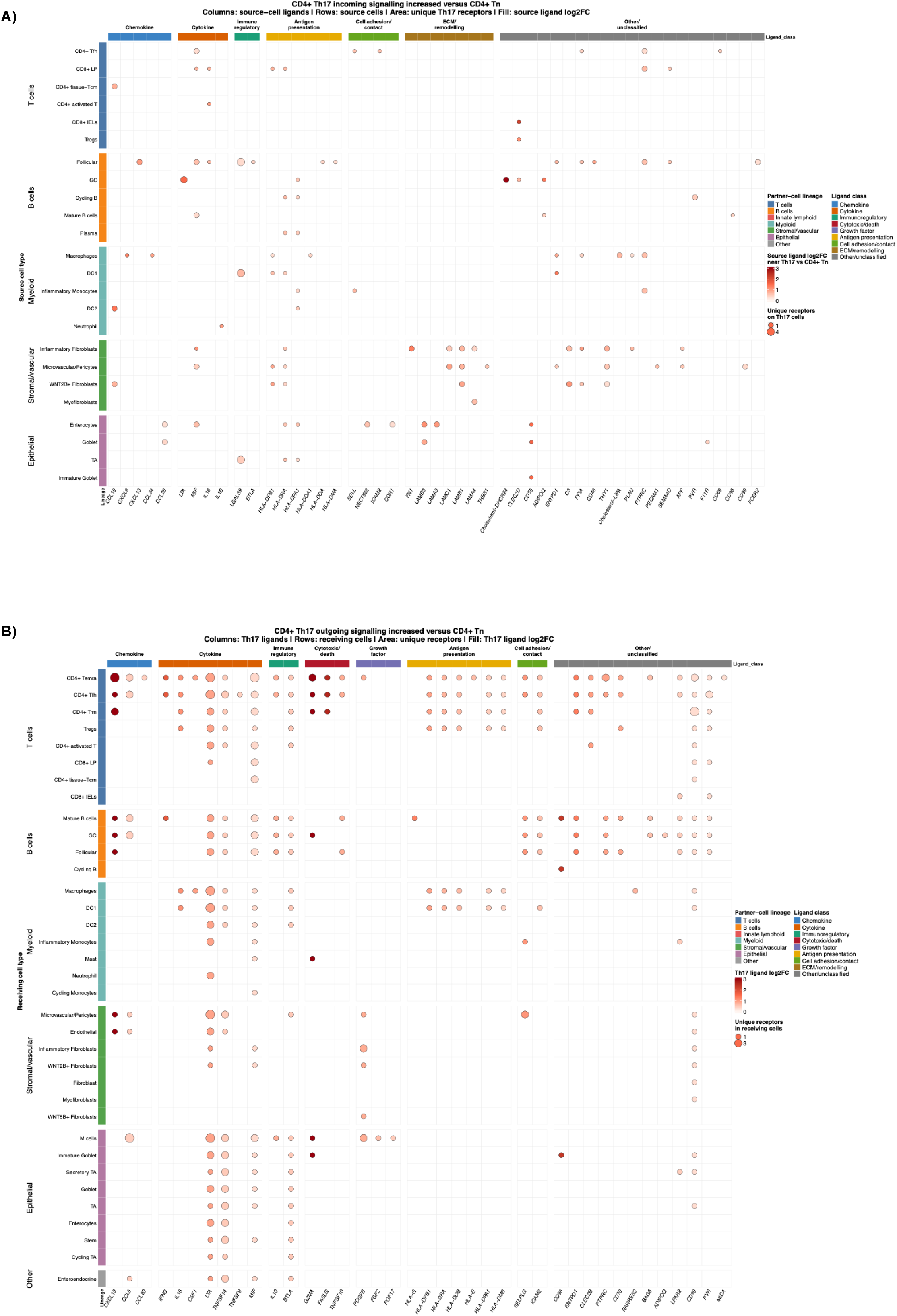
Differential spatial ligand-receptor interactions involving CD4⁺ Th17 cells relative to naïve CD4⁺ T cells. **A,** Candidate incoming signals received by CD4⁺ Th17 cells that were increased relative to CD4⁺ naïve T cells. Rows indicate spatially associated ligand-producing source populations and columns indicate source cell ligands, grouped by functional class. Dot size represents the number of unique receptors for the corresponding ligand detected on Th17 cells, while dot colour represents the log₂ fold change in source cell ligand expression in Th17-associated relative to naïve-T-cell-associated spatial neighbourhoods. **B,** Candidate outgoing signals from CD4⁺ Th17 cells that were increased relative to naïve CD4⁺ T cells. Rows indicate spatially associated receiving populations and columns indicate ligands expressed by Th17 cells, grouped by functional class. Dot size represents the number of unique corresponding receptors detected within each receiving population, while dot colour represents the log₂ fold change in ligand expression in Th17 relative to naïve CD4⁺ T cells. Coloured strips denote the major lineage of the interacting partner population, and column headers denote ligand functional class. Only interactions supported by significantly increased ligand expression at a Benjamini–Hochberg false-discovery rate <0.05 are shown. ECM, extracellular matrix; GC, germinal centre; IEL, intraepithelial lymphocyte; LP, lamina propria; TA, transit-amplifying cell; Tcm, central memory T cell; Temra, terminally differentiated effector memory T cell; Tfh, follicular helper T cell; Tn, naïve T cell; Trm, tissue-resident memory T cell.

**Supplementary Figure 6:**
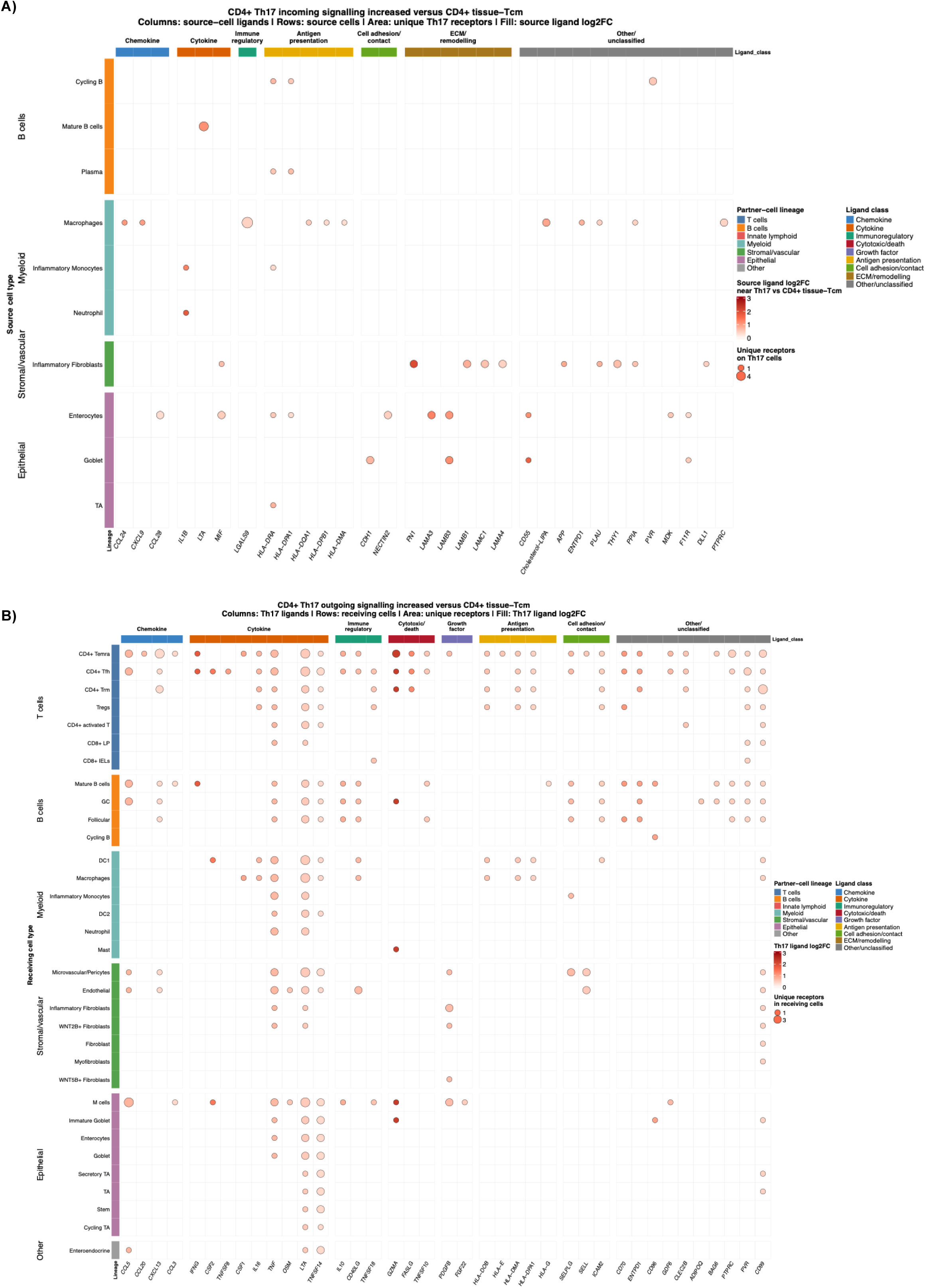
Differential spatial ligand–receptor interactions involving CD4⁺ Th17 cells relative to tissue central memory CD4⁺ T cells. **A,** Candidate incoming signals received by CD4⁺ Th17 cells that were increased relative to tissue central memory CD4⁺ T cells. Rows indicate spatially associated ligand-producing source populations and columns indicate source-cell ligands, grouped by functional class. Dot size represents the number of unique receptors for the corresponding ligand detected on Th17 cells, while dot colour represents the log₂ fold change in source-cell ligand expression in Th17-associated relative to tissue-Tcm-associated spatial neighbourhoods. **B,** Candidate outgoing signals from CD4⁺ Th17 cells that were increased relative to tissue-Tcm cells. Rows indicate spatially associated receiving populations and columns indicate ligands expressed by Th17 cells, grouped by functional class. Dot size represents the number of unique corresponding receptors detected within each receiving population, while dot colour represents the log₂ fold change in ligand expression in Th17 relative to tissue-Tcm cells. Coloured strips denote the major lineage of the interacting partner population, and column headers denote ligand functional class. Only interactions supported by significantly increased ligand expression at a Benjamini-Hochberg false-discovery rate <0.05 are shown. ECM, extracellular matrix; GC, germinal centre; IEL, intraepithelial lymphocyte; LP, lamina propria; TA, transit-amplifying cell; Tcm, central memory T cell; Temra, terminally differentiated effector memory T cell; Tfh, follicular helper T cell; Trm, tissue-resident memory T cell.

**Supplementary Figure 7:**
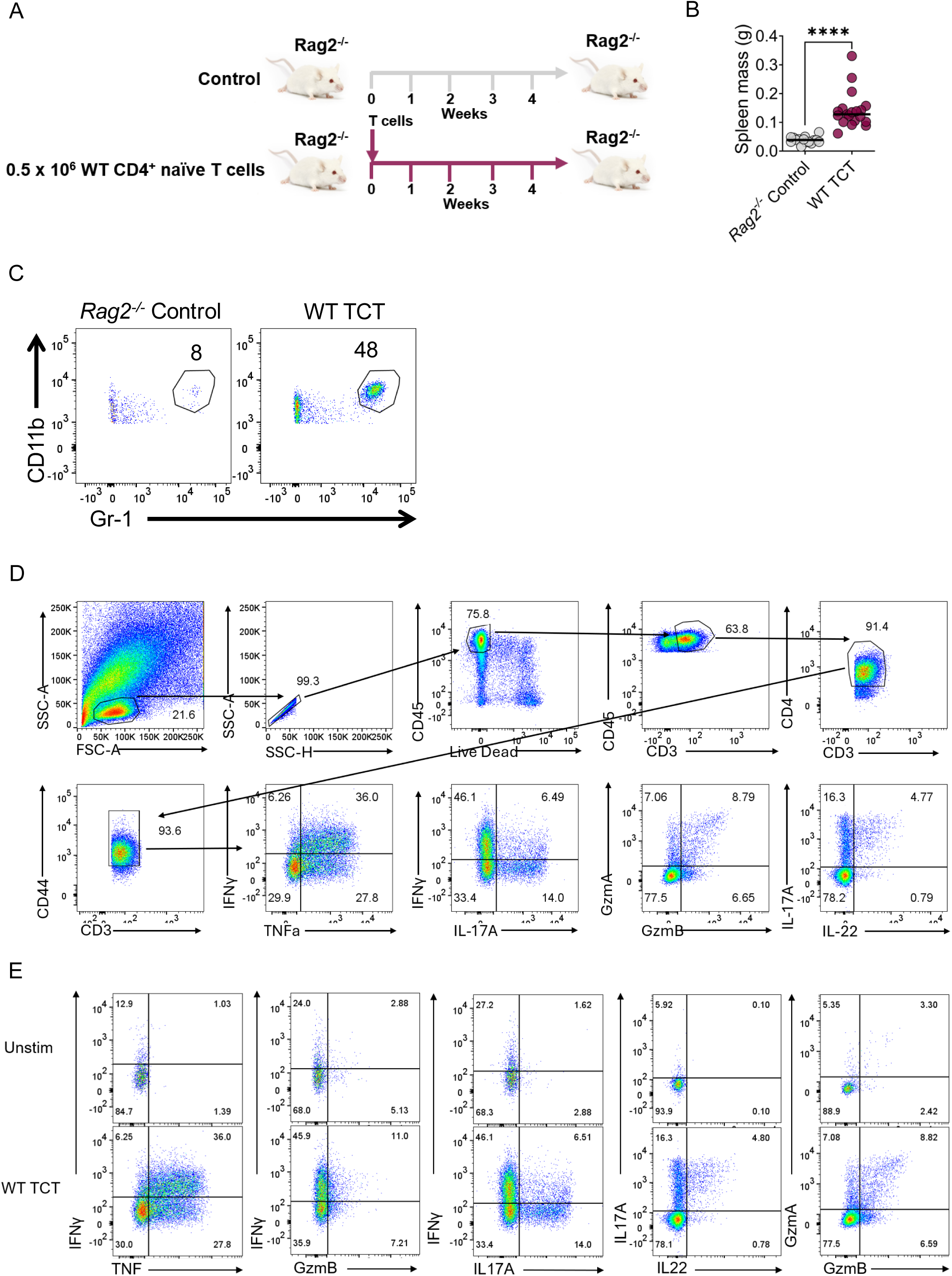
Wild-type CD4⁺ T-cell transfer induces colitis and a polyfunctional granzyme-expressing effector phenotype. **A,** Schematic of the CD4⁺ T-cell transfer colitis experiment. *Rag2*⁻/⁻ recipient mice received either no cells or 0.5 × 10⁶ WT naïve CD4⁺ T cells and were followed for four weeks. **B,** Spleen mass at the experimental endpoint. Each point represents one mouse; groups were compared using a two-sided Mann-Whitney U test. **C,** Representative flow cytometry identification of colonic CD11b⁺Gr-1^hi neutrophils in untreated *Rag2*⁻/⁻ controls and recipients of WT CD4⁺ T cells. Numbers indicate the percentages of cells within the displayed gates. **D,** Representative flow cytometry gating strategy used to identify viable colonic CD45⁺CD3⁺CD4⁺ donor-derived T cells and assess CD44, inflammatory cytokine and granzyme expression. Numbers indicate the percentages of cells within the corresponding gates or quadrants. **E,** Representative cytokine and granzyme co-expression plots for donor-derived CD4⁺ T cells recovered following WT T-cell transfer. Unstimulated controls used to establish quadrant gates are shown above the corresponding stimulated WT T-cell-transfer samples. \*\*\*\**P* < 0.0001. TCT, T-cell transfer; WT, wild type.

**Supplementary Figure 8:**
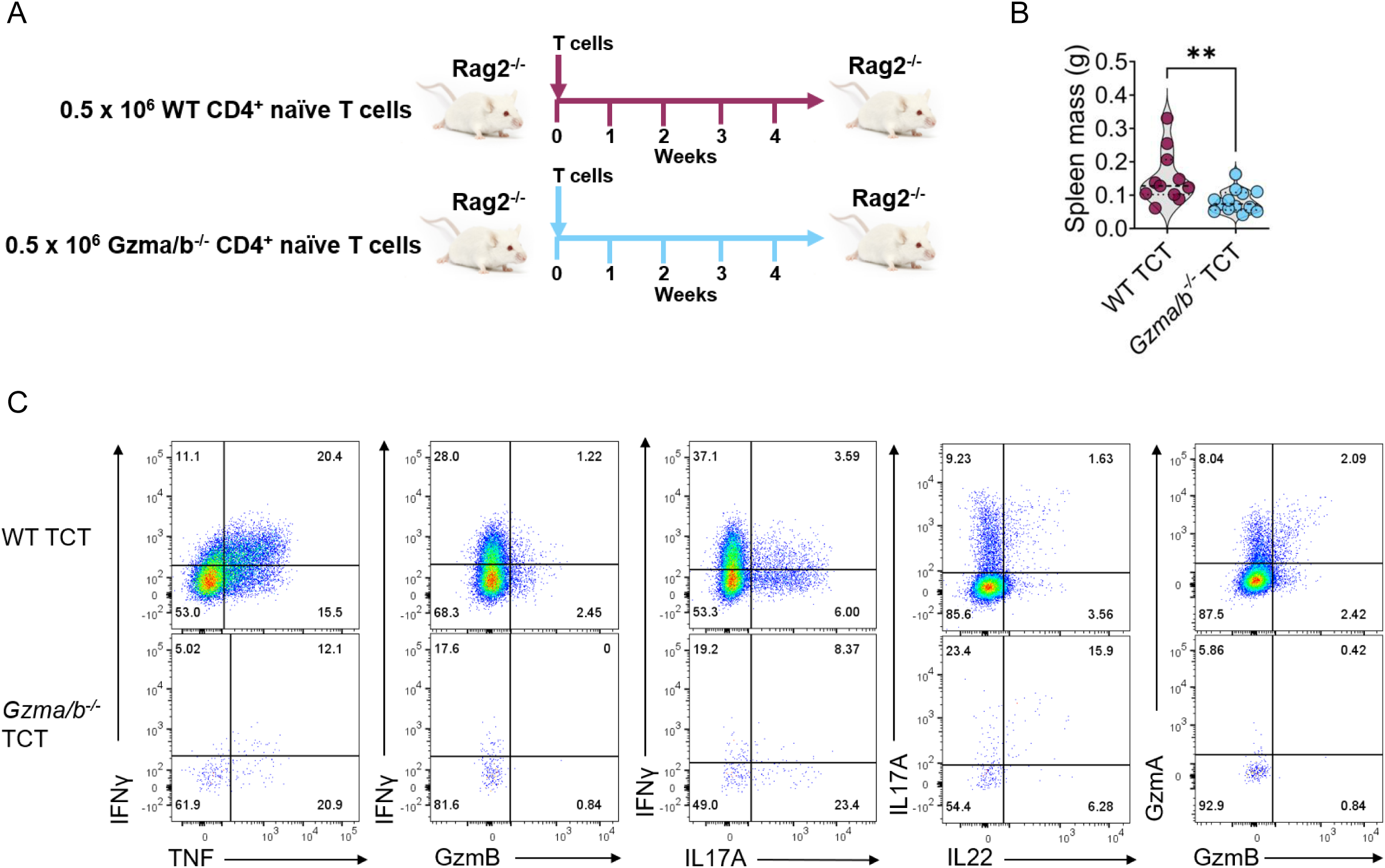
Transfer of GZMA/GZMB-deficient CD4⁺ T cells attenuates systemic inflammation and abrogates the polyfunctional granzyme-expressing phenotype. **A,** Schematic of the comparative CD4⁺ T-cell transfer experiment. *Rag2*⁻/⁻ recipient mice received 0.5 × 10⁶ WT or *Gzma/Gzmb*-deficient naïve CD4⁺ T cells and were followed for four weeks. **B,** Spleen mass at the experimental endpoint. Each point represents one mouse and groups were compared using a two-sided Mann–Whitney U test. **C,** Representative flow-cytometric comparison of inflammatory cytokine and granzyme co-expression among donor-derived colonic CD4⁺ T cells recovered following WT or *Gzma/Gzmb*-deficient T-cell transfer. Numbers indicate the percentages of cells within the corresponding quadrants. \*\**P* < 0.01. TCT, T-cell transfer; WT, wild type.

---

**Supplementary Table 1:** Markers included in the multiparameter flow-cytometry panel used to characterise conventional and unconventional T-cell populations in UC patients and non-IDB controls.

| Category | Marker |
| --- | --- |
| Cytotoxic molecule | GZMB |
| Cytokines | IFN-γ, IL-17A, IL-22, TNF-α |
| Conventional T cell | CD3, CD4, CD8 |
| Innate-like / unconventional T cell | CD161, CD56, TCR Vα24-Jα18 (iNKT), TCR $\gamma/\delta$, TCR Vα7.2 (MAIT) |
| Pan-leucocyte | CD45 |
| Live-dead | A700 |

**Supplementary Table 2:** Demographic and clinical characteristics of patients with active UC included in the spatial transcriptomics cohort. Data are presented as median (interquartile range) for continuous variables and *n* (%) for categorical variables. E1, proctitis; E2, left-sided colitis; E3, extensive colitis; SCCAI, Simple Clinical Colitis Activity Index; UC, ulcerative colitis.

| Characteristic | Active UC cohort (N = 16) |
| --- | --- |
| Age at endoscopy, years | 34.5 (27.0-50.8) |
| Female sex | 16 (100%) |
| Disease duration, years | 5.0 (2.0–21.5) |
| Disease extent |  |
| — Proctitis (E1) | 1 (6.3%) |
| — Left-sided colitis (E2) | 11 (68.8%) |
| — Extensive colitis (E3) | 4 (25.0%) |
| Biologic-naïve | 12 (75.0%) |
| Endoscopically active disease | 16 (100%) |
| SCCAI | 6.5 (3.5-8.3) |

**Supplementary Table 3:** Relative enrichment of the polyfunctional cytotoxic CD4⁺ T-cell signature across spatially resolved CD4⁺ T-cell subsets. Each CD4⁺ T-cell subset was compared with the pooled remaining CD4⁺ T-cell populations using a linear mixed-effects model incorporating participant identity as a random intercept. Positive estimates indicate greater signature enrichment relative to the remaining populations. *P* values were adjusted across subtype comparisons using the Benjamini-Hochberg procedure. CI, confidence interval; Tcm, central memory T cell; Temra, terminally differentiated effector memory T cell; Tfh, follicular helper T cell; Trm, tissue-resident memory T cell.

| CD4<sup>+</sup> T-cell subtype | Estimated difference in enrichment versus remaining CD4 T-cell subtypes | 95% CI | Adjusted p-value |
| --- | --- | --- | --- |
| Activated T cells | −0.065 | −0.093 to −0.038 | <0.0001 |
| Temra cells | 0.095 | 0.058 to 0.132 | <0.0001 |
| Tfh cells | −0.019 | −0.045 to 0.007 | 0.164 |
| Th17 cells | 0.149 | 0.121 to 0.177 | <0.0001 |
| Tissue-Tcm cells | −0.061 | −0.087 to −0.035 | <0.0001 |
| Naïve T cells | −0.104 | −0.130 to −0.077 | <0.0001 |
| Trm cells | 0.005 | −0.026 to 0.036 | 0.744 |

**Supplementary Table 4:** Primary and UC-restricted sensitivity analyses of the association between stool dysbiosis and mucosal polyfunctional cytotoxic CD4⁺ T-cell signature enrichment. Standardised β coefficients were estimated using multivariable linear regression adjusted for diagnosis, age, sex, biopsy location, Modified Baron score and the stool-biopsy collection interval, with the diagnosis term omitted from the UC-restricted analysis. Confidence intervals and two-sided *P* values were calculated using participant-clustered CR2 variance estimation with Satterthwaite-adjusted degrees of freedom. CI, confidence interval; UC, ulcerative colitis.

| Analysis | Samples | Participants | Standardised $\beta$<br>for dysbiosis | 95% CI | p-value |
| --- | --- | --- | --- | --- | --- |
| Primary: UC<br>and non-IBD<br>colonic<br>samples | 77 | 46 | 0.233 | 0.022 to 0.444 | 0.033 |
| Sensitivity:<br>UC-only<br>colonic<br>samples | 49 | 25 | 0.308 | -0.047 to 0.663 | 0.082 |

